# Constructing the Human Brain Metabolic Connectome Using MR Spectroscopic Imaging: Insights into Biochemical Organization, Cytoarchitectonic Similarity and Gene Co-expression Networks

**DOI:** 10.1101/2025.03.10.642332

**Authors:** Federico Lucchetti, Edgar Céléreau, Pascal Steullet, Yasser Alemán-Gómez, Patric Hagmann, Antoine Klauser, Paul Klauser

## Abstract

Network science has revolutionized our understanding of brain organization by revealing self-organizing patterns underlying its structural and functional connectivity. However, capturing metabolic contrast remains a challenge, leaving a critical gap in connectomics. Using advanced 3D whole-brain proton MR spectroscopic imaging with high spatial resolution and shortened acquisition times, we constructed the first human brain metabolic connectome in 51 healthy subjects, validated on an independent cohort (N=12) scanned at a different site. Our pipeline generates consistent and reliable metabolic similarity matrices. Upon further analysis, metabolic similarity networks display distinct topological features, notably a smoothly varying gradient delineating functionally and spatially distinct yet integrated brain regions via connector hubs. Although metabolic hubs correlate with structural hubs, overall alignment with structural connectivity is poor. However, metabolic organization aligns more closely with cytoarchitectonic and genetic co-expression patterns, suggesting a neurodevelopmental origin. This work puts forward the metabolic similarity gradient as a hallmark of the brain’s overarching biochemical organization, and provides a foundation for incorporating metabolite imaging into the broader domain of connectomics and its potential applications in health and disease.

## 1 Introduction

Since their inception in the 1980s, magnetic resonance imaging (MRI) techniques have become central to neuroimaging. MRI offers non-invasive, non-ionizing measurements of the living brain across various ages, with relatively quick acquisition times and moderate costs, making it widely accessible. These advantages have established MRI as a vital tool in both clinical and basic neuroscience.

MRI leverages the resonance properties of ^1^H nuclear spins to create contrasts based on relaxation (e.g., T1-, T2-, and T2*-weighted imaging like BOLD fMRI), diffusion (e.g., diffusion tensor imaging), and chemical shifts (e.g., magnetic resonance spectroscopy). These techniques have not only enhanced our understanding of the brain’s anatomical structure, composition, and function but also enabled the construction of large-scale brain network models.

In particular, diffusion-weighted imaging (DWI) maps water diffusion along neuronal axons, revealing white matter tracts and pioneering the structural connectome and the field of connectomics Hagmann (2005); Sporns, Tononi, and Kötter (2005). BOLD signals add a temporal dimension, defining functional connectivity as the statistical relationships between regional brain activities, such as correlated blood flow or synchronized neural signals.

The field of connectomics has revealed that, like many other complex networks, both structural and functional brain networks exhibit fundamental properties, including a modular organization that balances segregation and integration Bullmore and Sporns (2012); Chen, Wang, Hilgetag, and Zhou (2017); Ma et al. (2023). Local circuits form dense, spatially segregated communities that optimize metabolic costs and support specialized cognitive functions, sensory processing, motor control, and other localized activities. Meanwhile, long-range connections, despite their metabolic expense, enable integration through connector hubs or rich-club nodes, facilitating efficient communication between distant and functionally distinct regions. Individual differences in brain network organization are linked to variations in behavior, including general intelligence, working memory, and personality traits, and have significant implications for understanding neurological and psychiatric disorders van den Heuvel and Sporns (2019). Referred to as connectopathies Martin (2012), these disorders are increasingly recognized as arising from disruptions within large, distributed brain networks Bullmore and Sporns (2009); de Lange et al. (2019); Stam (2014); Xia, Sun, Bu, Li, and He (2024). Network abnormalities often center on inter-hub connector nodes, which incur higher metabolic costs in terms of oxidative glucose metabolism Vaishnavi et al. (2010) and become vulnerable to homeostatic disturbances. Consequently, they represent potential failure points in various pathological conditions Bullmore and Sporns (2012); de Lange et al. (2019). In vivo diffusion MRI captures macro-circuit changes (e.g., fractional anisotropy, tractography) Steullet et al. (2016), yet detecting microcircuit alterations remains challenging with structural MRI alone. Functional MRI and dynamic causal modeling of functional networks Friston (2002); Stephan et al. (2008) provide insights into metabolic activity Seghier, Zeidman, Neufeld, Leff, and Price (2010), but remain limited by computational intensity, model specificity, and interpretative complexity Heinzle and Stephan (2018).

Thus given that structural and functional networks are likely underpinned by the brain’s metabolic organization, their limitation to capture directly metabolic activity has left a critical gap in connectomics, underscoring the timely need for new imaging methods capable of rendering the brain’s metabolic connectome.

In this paper, we propose leveraging recent advancements in proton magnetic resonance spectroscopic imaging techniques Klauser et al. (2019); Klauser, Klauser, Grouiller, Courvoisier, and Lazeyras (2022); Klauser, Strasser, Thapa, Lazeyras, and Andronesi (2021); Lecocq et al. (2015); Maudsley et al. (2009); Moser et al. (2020); Steel et al. (2018), which enable whole-brain 3D metabolite imaging at high resolution with significant reduced acquisition times. Building on the principles of MRS, which suppresses water and lipid signals to non-invasively quantify metabolic compounds in the MR spectrum, including the five main spectrally resolved metabolite contents : N-acetylaspartate + N-acetyl aspartylglutamate (tNAA), creatine + phosphocreatine (tCr), glutamate + glutamine (Glx), phosphocholine + glycerophosphocholine (Cho), and myo-inositol (Ins), ^1^H-MRSI facilitates detailed, scalable analyses of brain metabolism. This technique can be easily integrated into an MRI protocol for clinical or research purposes and, to the best of our knowledge, enables the construction of the first human brain MRSI-based metabolic connectome, capturing metabolic correlations across brain regions, which is the core focus of this study.

We explore the feasibility of constructing an individual metabolic similarity matrix (MeSiM) using data from a single whole-brain scan ^1^H-MRSI in a cohort of healthy participants. We test four central hypotheses regarding its network structure: (1) its topological characteristics mirror those of naturally occurring networks; (2) it exhibits a modular organization; (3) it correlates with structural connectivity, cytoarchitectonic similarity, and genetic co-expression patterns; and (4) regions with the highest structural connectivity coincide with those of high metabolic concentration.

## 2 Results

### 2.1 Metabolic Similarity Matrices

#### 2.1.1 Construction

To evaluate the feasibility of a within-subject metabolic similarity matrix (MeSiM), we analyzed ^1^H-MRSI data from 51 healthy adolescents in Geneva (i.e., Mindfulteen study Piguet et al. (2022)) following the similarity matrix construction pipeline schematically laid out on Fig. 1. Each anatomical volume was parcellated using the LFMIHIFIF-3 atlas, which defines 277 brain regions (Methods 7.2.1). Largely following the same methodology used to extract morphometric profiles Seidlitz et al. (2018) and detailed in Methods 7.2.5), we extracted, based on five spectrally resolved proton-MRSI metabolic compounds (tNAA, tCr, Glx, Cho, Ins) , and retained 210 z-score-normalized metabolic profiles corresponding to 210 brain regions with reliable ^1^H-MRSI coverage (see Extended Data 8.1).

**Fig. 1.**
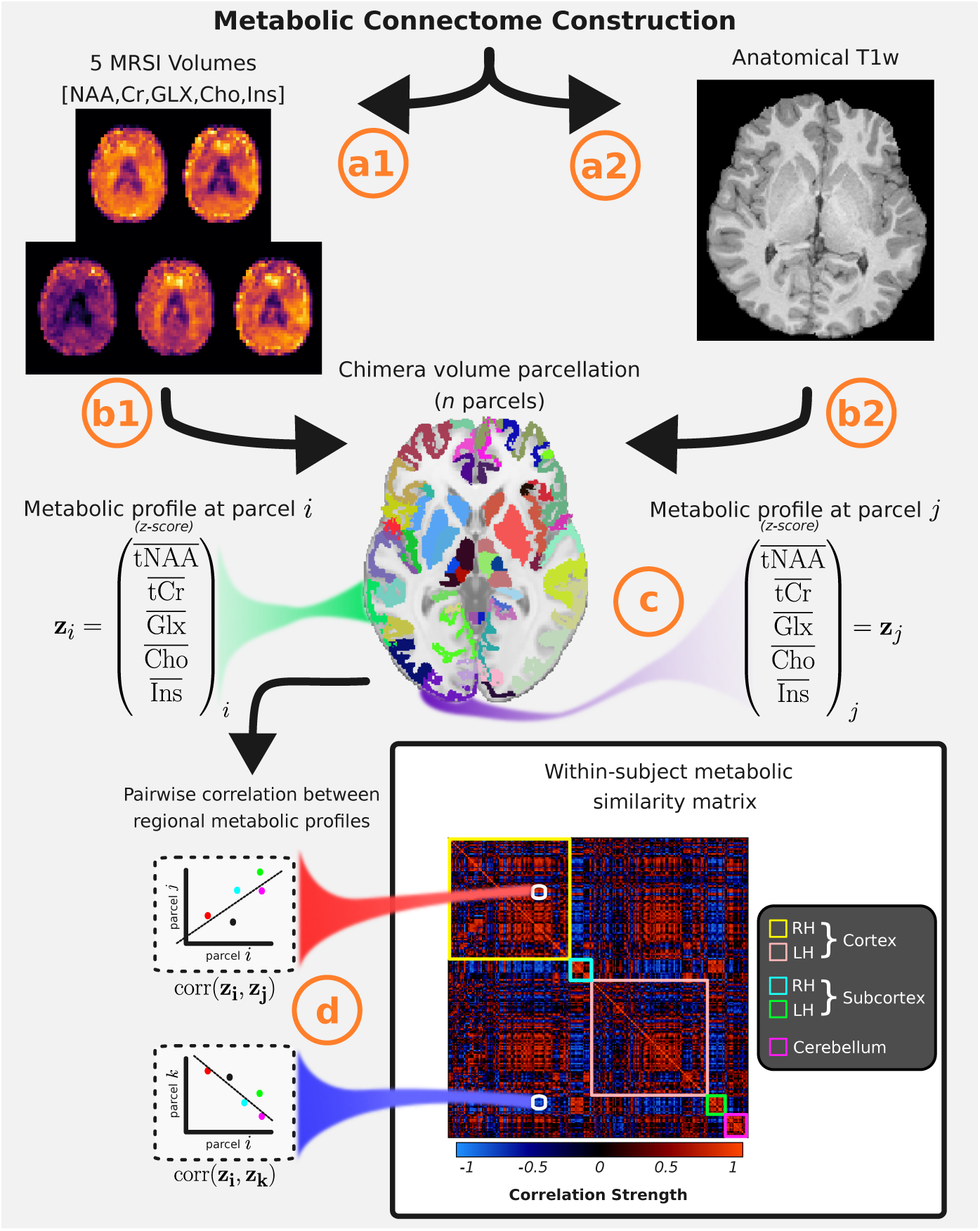
Individual Metabolic Similarity Matrix Construction. **a1** & **a2**, Acquisition of ^1^H-MRSI and anatomical T1-weighted images, followed by the reconstruction of 5 spectrally resolved metabolite brain concentrations, resulting in 5 volume maps ([*tNAA, tCr, Glx, Cho, Ins*]). **b1**. Coregistration of the anatomical image to the ^1^H-MRSI space and mapping of the five metabolite concentrations to the *n* brain parcels. **b2**, Anatomical volume parcellation using the Chimera parcellation scheme Alemán-Gómez (2024), producing *n* brain parcels. **c**, Calculation of the median metabolite concentration per parcel, followed by z-score normalization across all brain parcels per metabolite for each individual, to generate standardized metabolic profiles *{***z***_m_}_m_*_=1,*…,n*_, where *{***z***_i_}* denotes the z-score normalized metabolic profile at region *i*. **d**, Construction of pairwise Spearman correlations between metabolic feature vectors across all brain regions, yielding a within-subject metabolic similarity matrix (MeSiM).

The metabolic profiles were augmented via a perturbation procedure (see Methods 7.2.5), increasing the sample size per metabolite concentration by a factor of *K_pert_*. Pairwise Spearman correlations between brain regions were calculated to create a 210 × 210 MeSiM for each individual. Fig. 2a shows an example MeSiM for *K_pert_* = 50, highlighting strong positive correlations clustering along the diagonal, representing super-regional communities (cortex, subcortex, cerebellum). Symmetrical clusters between hemispheres indicate metabolic similarity in corresponding regions, while negative correlations off-diagonally, particularly between distinct super-regions (e.g., cortex and subcortex), reflect metabolic dissimilarity and are likely driven by differences in tissue composition.

**Fig. 2.**
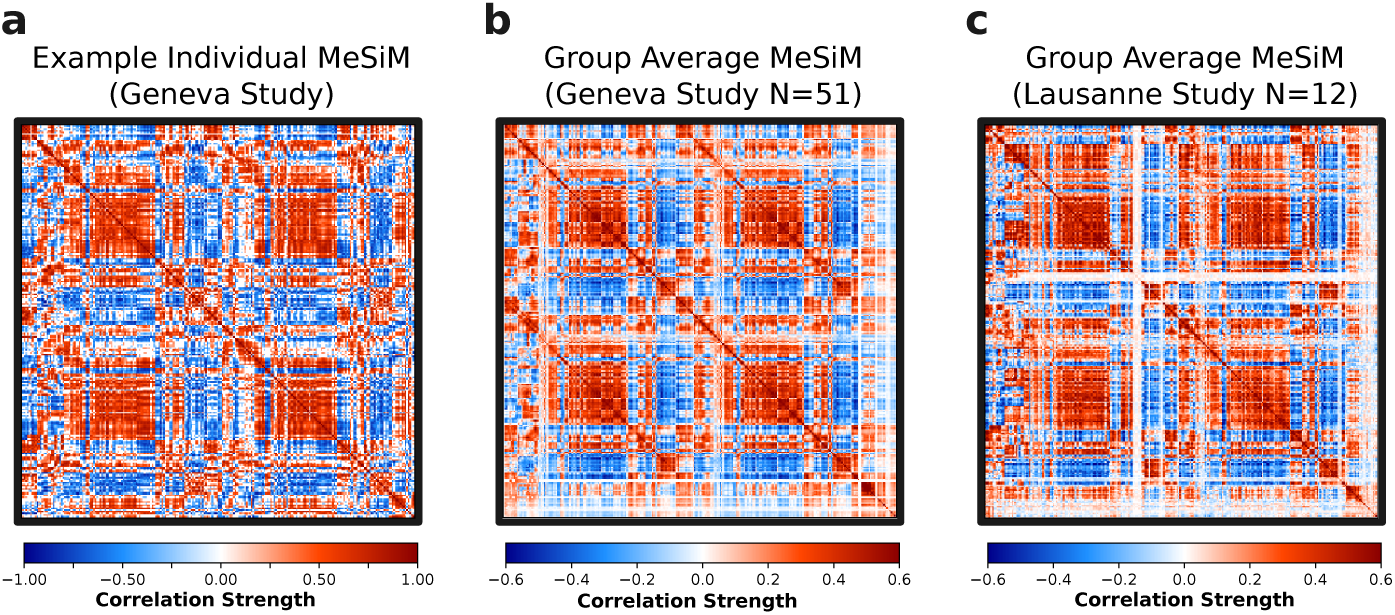
Individual and Group Average Metabolic Similarity Matrices. **a**, Individual MeSiM (*N* = 1), from the Geneva study **b**, Group averaged (*N* = 51) MeSiM from the Geneva study. **c**, Group averaged (*N* = 12) MeSiM from the Lausanne study.

#### 2.1.2 Stability, Consistency and Replicability

We evaluated the stability of MeSiM construction by testing two key parameters: the role of individual metabolites and *K*_pert_ values. A leave-one-metabolite-out approach assessed the influence of individual metabolites by comparing the original 5-metabolite MeSiM with MeSiM matrices reconstructed after imputing a single metabolite. This was repeated for three *K*_pert_ values (1, 50, 100) in all individuals. For each test, pairwise edge strengths from the original and imputed matrices were correlated for each individual’s MeSiM, and these correlations were aggregated across all subjects using Fisher’s z-transformation, with p-values corrected using the Benjamini-Hochberg false discovery rate method. At *K*_pert_ = 1, correlations between the original and leave-one-out MeSiM were low, ranging from *r* = 0.20 to *r* = 0.29 (*p <* 0.05). Increasing *N*_pert_ to 50 significantly improved these correlations (tNAA (*r* = 0.79), Ins (*r* = 0.96), Cho (*r* = 0.81), Glx (*r* = 0.84) and tCr (*r* = 0.95), all *p <* 0.05), with marginal gains at *K*_pert_ = 100 (*r* = 0.80 to *r* = 0.96, *p <* 0.05). To further examine the differences in contribution between metabolites, an ANOVA test was performed on the correlations between leave-one metabolite-out at *K_pert_* = 50 revealing a significant difference between metabolites (*F* = 1154, *p <* 0.001). A post hoc Tukey HSD test indicated that tNAA, Cho, and Glx contributed significantly more than the other metabolites to the MeSiM construction. In contrast, removing Ins or tCr had minimal effects on correlation values. The consistency of MeSiMs was evaluated by comparing individual MeSiMs with a group-level MeSiM, derived by averaging all individual MeSiMs from the Geneva study (Fig. 2b). Variability was quantified by calculating edge-wise correlations between individual- and group-level MeSiMs, Fisher-transformed, and aggregated for overall alignment. Correlations increased from *r* = 0.33 at *K*_pert_ = 1 to *r* = 0.55 at *K*_pert_ = 50, with a minor improvement at *N*_pert_ = 100 (*r* = 0.56), showing that *K*_pert_ = 50 sufficiently reduces variability and ensures consistent MeSiM construction. We fixed *K*_pert_ = 50 for subsequent analyses. Replicability was assessed using a different sample of healthy participants from Lausanne (i.e., Lausanne Psychosis Cohort Baumann et al. (2013) ) scanned on a different MRI platform. The group-level MeSiMs for the Geneva and Lausanne studies (shown on Fig. 2c) were strongly correlated (*r* = 0.86, *p <* 0.001), demonstrating the replicability of the MeSiM construction method across sites (i.e., different sample and scanner).

### 2.2 Metabolic Similarity Gradient

To show the spatial organization of metabolic similarity and the spatial discrimination between positive and negative MeSiM weights, we analyzed pairwise group-averaged MeSiM weights (*N* =51, Geneva study) in relation to inter-regional Euclidean distances, pooling data from both hemispheres (Fig. 3a). We then binned positive and negative MeSiM values separately in 2 mm intervals, generating density distributions (Fig. 3b). Positve MeSiM values cluster at shorter distance ranges (37mm), whereas negative MeSiM values peak at longer distance ranges (71 mm). We fitted these distributions with lognormal models and evaluated goodness-of-fit (GoF) using spatially informed null models to generate null distributions. Given the distinct spatial roles of positive and negative MeSiM, we applied two separate parcel permutation procedures (Methods 7.4.2). The adjacent parcel permutation test (AdjPerm) generated a null distribution by permuting MeSiM values only between adjacent parcels, thereby preserving local structure and testing the significance of the positive MeSiM value fit against the chance distribution. The volumetric spin test (VolSpin) generated a null distribution by performing multiple solid rotations of cortical and subcortical regions along two concentric spheres, preserving large-scale gradients and anatomical spatial relationships to determine whether the observed spatial pattern of negative MeSiM values differs meaningfully from chance. Positive MeSiM weights showed GoF = 0.93 (AdjPerm *p* = 0.038), with a mode at 37 mm, negative MeSiM weights showed GoF = 0.97 (VolSpin *p <* 0.001), with a mode at 71 mm. The lognormal distribution for positive weights aligns with parcel spatial boundaries, exhibiting a sharp rise below 44 mm due to spatial constraints (average parcel radius ≈ 22 mm), followed by a decline as metabolic similarity decreases with distance. These findings suggest that metabolically similar regions shape local structure, whereas dissimilar regions define global topology, reinforcing the biological plausibility of metabolic connectivity and the fundamental role of spatial constraints in brain network economy Bullmore and Sporns (2012). To clearly visualize its spatial embedding, we reduced the 210×210 group-averaged MeSiM matrix to a 210×1 feature vector using Principal Component Analysis (PCA) for denoising and variance capture, followed by t-distributed Stochastic Neighbor Embedding (t-SNE) to preserve the local connectivity patterns in the MeSiM matrix (see Methods 7.2.6). The resulting 210×1 representation, termed the Metabolic Similarity Index (MSI), was projected back to the geometric locations of the corresponding brain parcels in MNI space to enable a 3D visualization of metabolic similarity patterns (Fig. 3c). The MSI values were inverse-mapped to the five z-score-normalized metabolic concentration profiles ( Fig. 3c attached to color scale), allowing each MSI value to be interpreted as a reduced representation of the 5-feature metabolic profile. This representation reveals a bilateral symmetry, spatial clustering of metabolically similar regions, and a clear separation between cortical and subcortical regions. The latter is largely driven by the flipping tNAA-to-Cho ratio (tNAA dominating in cortical regions and Cho in subcortical regions), which underpins the observed inversion of predominant positive MeSiM values (cortical-to-cortical connections) to negative ones (cortical-to-subcortical connections). The MSI gradient captures smooth variations within the cortex following a rostro-caudal path and delineates distinct brain regions such as the primary motor cortex (ie, precentral gyrus) and the primary somatosensory cortex (ie, postcentral gyrus). The limbic system, comprising the insula, cingulate cortex and other subcortical structures (i.e., basal ganglia, thalamus, hypothalamus and hippocampus) show similar metabolic pattern clearly distinguishable from the isocortex. Interestingly, the rostro-caudal gradient also appears within well-defined structures such as the cingulate cortex or insula. The cerebellum metabolically aligns with the subcortical-prefrontal axis (yellow-to-red MSI) but remains distinct from the nearby occipital lobe. This indicates that the spatial autocorrelational structure of MeSiM weights is likely due to biological processes rather than methodological artifacts (e.g., spatial blurring). Additionally, MSI map generation is robust to cortical parcellation (see Extended Data 8.3).

**Fig. 3.**
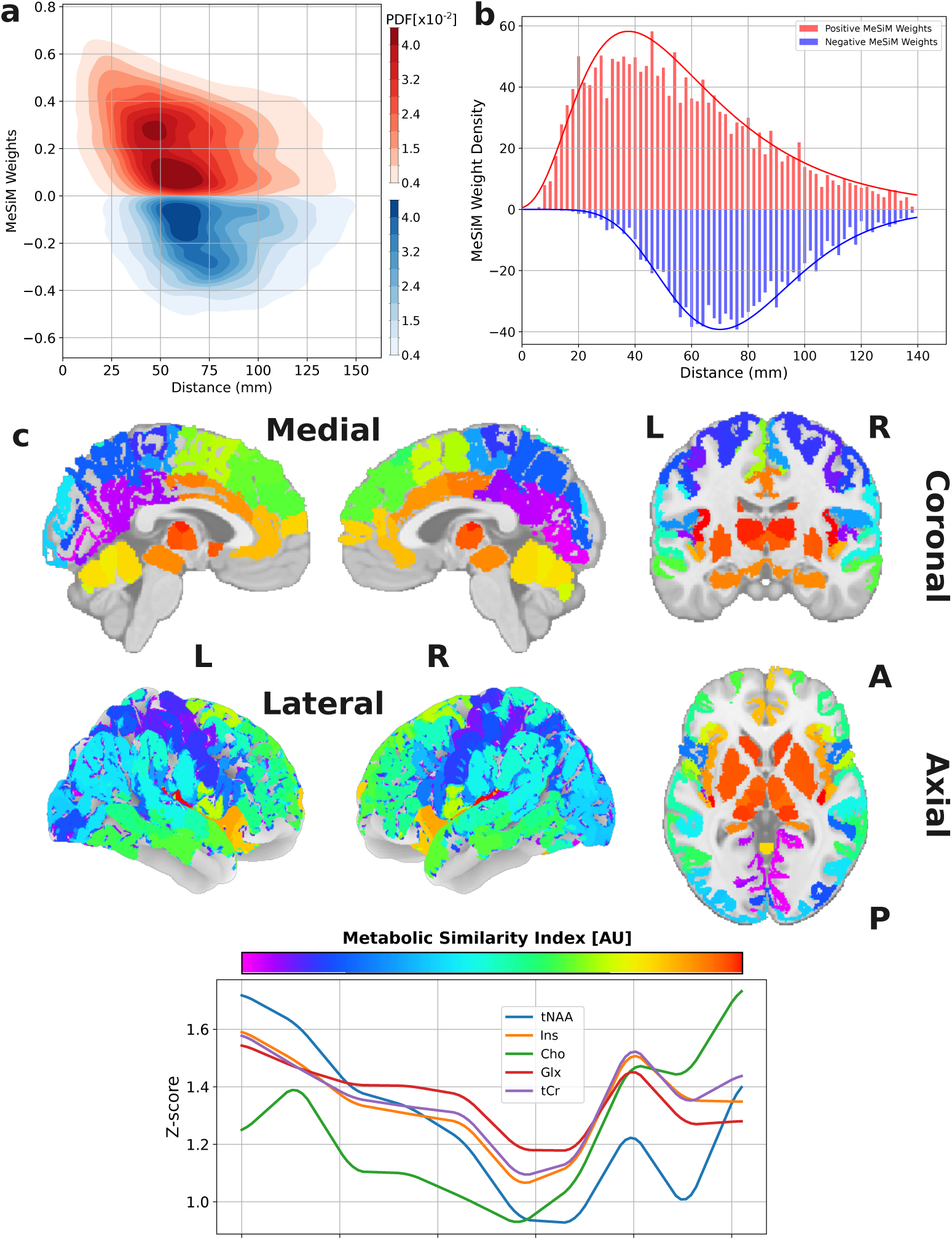
Metabolic Scalar Index. **a**, Density of negative (blue) and positive (red) metabolic pairwise correlations pooled from the group averaged (*N* = 51) MeSiM Geneva study aggregated across both hemispheres as a function of their respective inter-region Euclidean distance. Histogram values (z-axis) were smoothed using a kernel density estimation technique with Gaussian kernels. **b**, Aggregated MeSiM weights summed within distance bins of 2 mm. Continuous lines represent a lognormal fit **c**, Spatial mapping of the group-averaged MeSiM matrix (Geneva study), reduced to a single component using PCA-t-SNE, projected onto MNI space, and represented by the Metabolic Similarity Index (MSI). The MSI is color-coded (color bar) and inverse-mapped to the five z-score-normalized metabolic concentration profiles.

### 2.3 Network Topology

In order to reveal metabolic similarity network topology, we thresholded individual MeSiMs (see Methods 7.5.1) to generate binary graphs (see left Fig. 4a for group level MeSiM binarization at 18% connection density) and calculated graph metrics across connection densities (2%, 3%, 4%, 5%, 10%, 20%, and 30%). At all densities, metabolic networks exhibited complex topological characteristics typical of naturally occurring networks Barabási (2013) with rich-club coefficients being statistically significant (*p <* 0.001) (see Methods 7.5.2 and Extended Data 8.4). These findings were replicated on the group-averaged MeSiM, and the resulting rich-club network is shown in Fig. 4a. The occipital lobe exhibited the highest density of rich-club nodes, with fewer observed in the frontal lobe. When coloring nodes and edges by their MSI, we highlight metabolically distinct and spatially segregated clusters, each containing at least one rich-club node (except the cerebellum). This underscores the natural community structure and the integration of metabolically similar yet spatially separated regions through highly interconnected nodes. We further highlighted the integrative nature of a subset of the rich-club nodes (connector hubs) by building on the previous observation that the MSI gradient traverses the brain’s main lobes (see top left panel on Fig. 4c) and whose curve coordinates can analytically be distilled using pathfinding methods analogous to tract tracing in diffusion MRI (see Methods 7.2.7). The resulting network paths were fitted with a piecewise cubic spline for visualization, 3D-rendered on Fig. 4b and schematically visualized in Fig. 4c. From the PCN, the primary curve begins in the visual pericalcarine region, traverses the lateral-inferior parietal lobe (IPL), and converges with a secondary curve originating in the superior parietal lobe (SPL) and precentral gyrus (PreC). It continues through the lateral frontal lobe and terminates at the rACC. Secondary branches extend from the IPL to the middle temporal gyrus (MTG) and from the rACC to the anterior insula (aIns) and medial prefrontal cortex (mPFC). A subcortical curve, structurally disjoint yet metabolically connected via an edge between the rACC and caudate nucleus (interrupted path), links the thalamus, brainstem hubs, and the cerebellum. Thus we identified curves aligned with the MSI gradient, termed *metabolic fibers* (similar to white matter tracts in diffusion MRI), which relay nodes through connector hubs, linking regions of metabolic similarity and defining the core topology of the metabolic connectome.

**Fig. 4.**
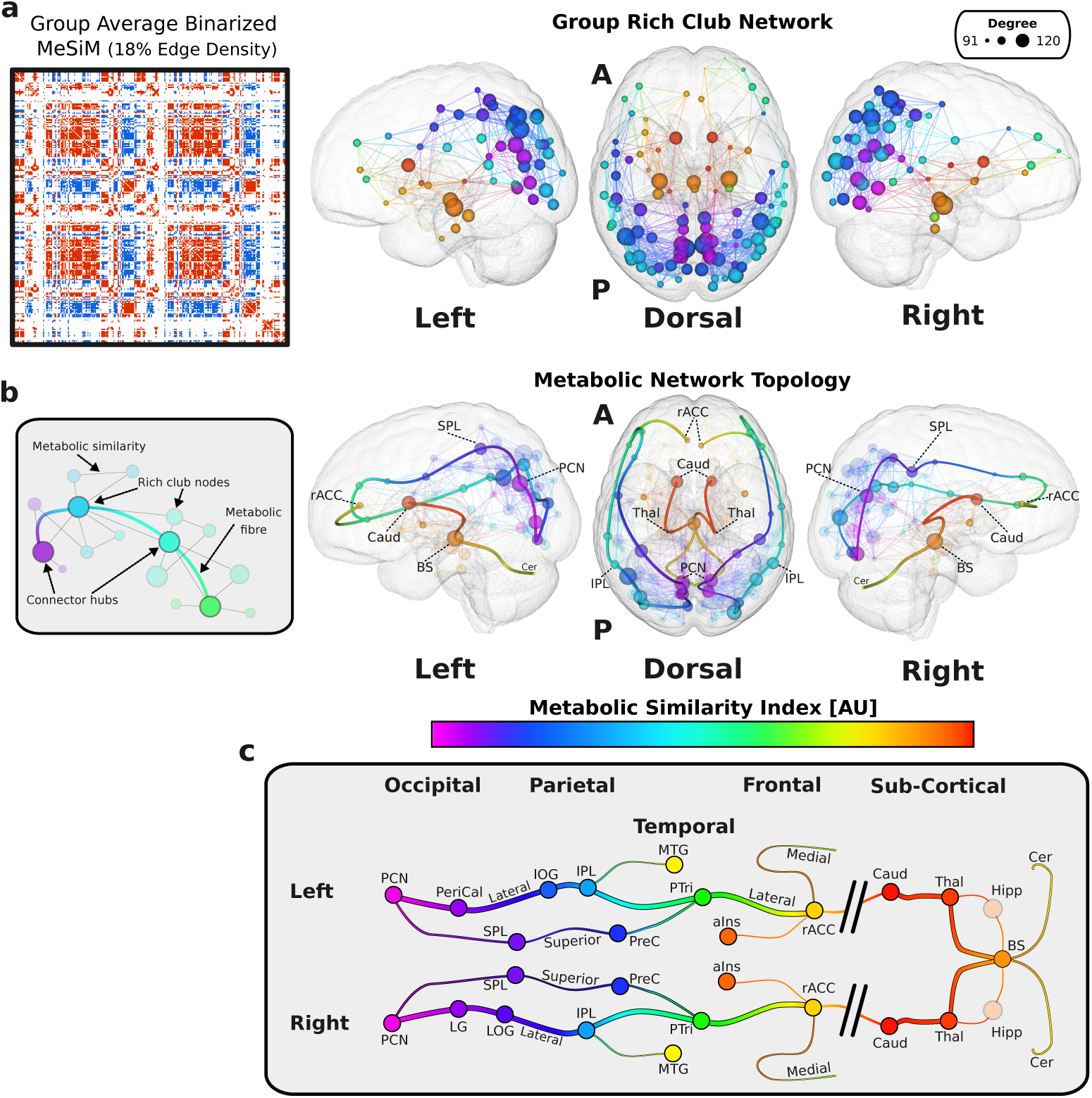
Metabolic Network Topology. **a**, Binarized Group MeSiM (18% edge density) from the Geneva study (left) and associated rich club network (right). Rich-club nodes are displayed at their anatomical locations, color-coded by their MSI, with node radii scaled according to degree. Interhemi-spheric edges are omitted for visualization purposes. **b**, Core topology of the metabolic connectome represented by connector hub situated along the metabolic fibers depicting regions of metabolic similarity through a smooth gradient color-coded by the MSI. Rich-club nodes (transparent) and edges are also highlighted. **c**, Schematic representation of the main metabolic fibers per hemisphere running through the cortex of four brain lobes and from subcortical structures to the brainstem and cerebellum, following the rostro-caudal axis. Connector hubs are highlighted along the curves. Metabolic similarity (but interrupted connection) between the subcortical fiber and the cortical fiber is indicated by a dashed curve. Precuneus (PCN), Pericalcarine cortex (PeriCal), Lingual gyrus (LG), Superior parietal lobule (SPL), Precentral gyrus (PreC), Lateral occipital gyrus (LOG), Inferior occipital gyrus (IOG), Inferior parietal lobule (IPL), Middle temporal gyrus (MTG), Pars triangularis (PTri), Rostral anterior cingulate cortex (rACC), Anterior insular cortex (aIns), Caudate nucleus (Caud), Thalamus (Thal), Brainstem (BS), Hippocampus (Hipp), and Cerebellum (Cer)

### 2.4 Comparative Analyses

#### 2.4.1 Metabolic Similarity and Structural Connectivity

To test the hypothesis that MeSiM edges reflect axonal connectivity, we analyzed diffusion spectrum imaging (DSI) data from the Geneva study, reconstructing tractograms and deriving weighted structural connectivity matrices (SC) based on the same Chimera gray matter parcellation LFMIHIFIF-3 (see Methods 7.2.2) to allow direct MeSiM and SC edge wise comparisons (see Fig. 5a). After excluding individuals with poor quality tractograms, 42 SC matrices were retained, with only the nodes covered by both matrices used to produce 42 190 × 190 matrices.

**Fig. 5.**
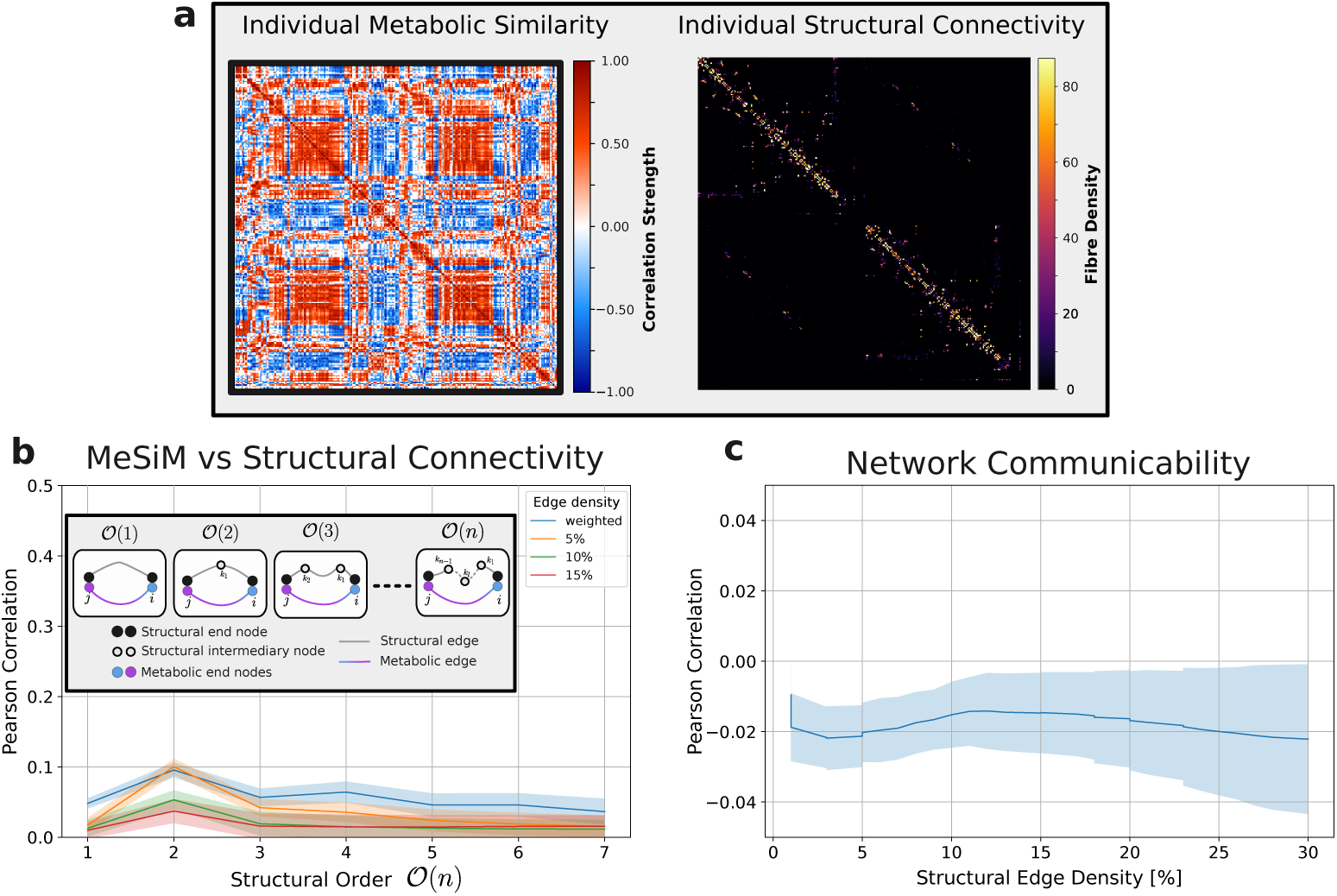
Metabolic Similarity and Structural Connectivity. **a**, Individual MeSiM and its corresponding structural connectivity matrix, represented as fiber density. **b**, Edge-wise correlation of structural connectivity matrices with MeSiM as a function of structural order. The orders *O*(1), *O*(2), *O*(3), and *O*(*n*) are schematically illustrated in the legend of (A_1_), where the *n*-th order involves *n −*1 intermediary structural nodes. **c**, Network communicability correlation (*M* = exp(*gS*)) between the metabolic similarity network (*M* ) and the structural connectivity network (*S*).

A direct edge-wise Pearson correlation between structural connectivity and metabolic similarity yielded a low scores (0.01 to 0.05, *p <* 0.001). We hypothesized that metabolic similarity between nodes could depend on indirect structural connections via intermediate nodes. By computing **S***^n^* (the *n*-th power of the SC matrix, see Methods 7.5.5), we found a peak correlation of 0.1 (*p <* 0.001) at second-order connections, which declined for higher orders (see Figure 5b). This suggests a modest influence of indirect structural pathways on metabolic similarity, with a weak overall relationship.

We also tested whether perturbations of metabolic nodes could be influenced by distant structural nodes using communicability analysis Estrada and Hatano (2008). Fig. 5c shows correlations between the predicted communicability model and the observed metabolic network were consistently weak (*r <* −0.022, *p <* 0.001) in binary and weighted matrices. These findings indicate that the metabolic similarity network is effectively independent of structural connectivity under the tested communicability model.

Lastly, we tested the hypothesis proposed by Hagmann et al. (2008); Vaishnavi et al. (2010) that structurally central nodes exhibit higher metabolic activity. Spearman correlations between structural degree centrality and metabolic profiles (z-score normalized) (sample correlation shown in Fig. 6a) yielded values ranging from 0.19 to 0.25 across connection densities (1%-20%) (see Fig. 6b). Taken together, these results confirm the poor resemblance between structural connectivity and metabolic similarity but establish a link between structural centrality and increased ^1^H-MRSI metabolite concentrations.

**Fig. 6.**
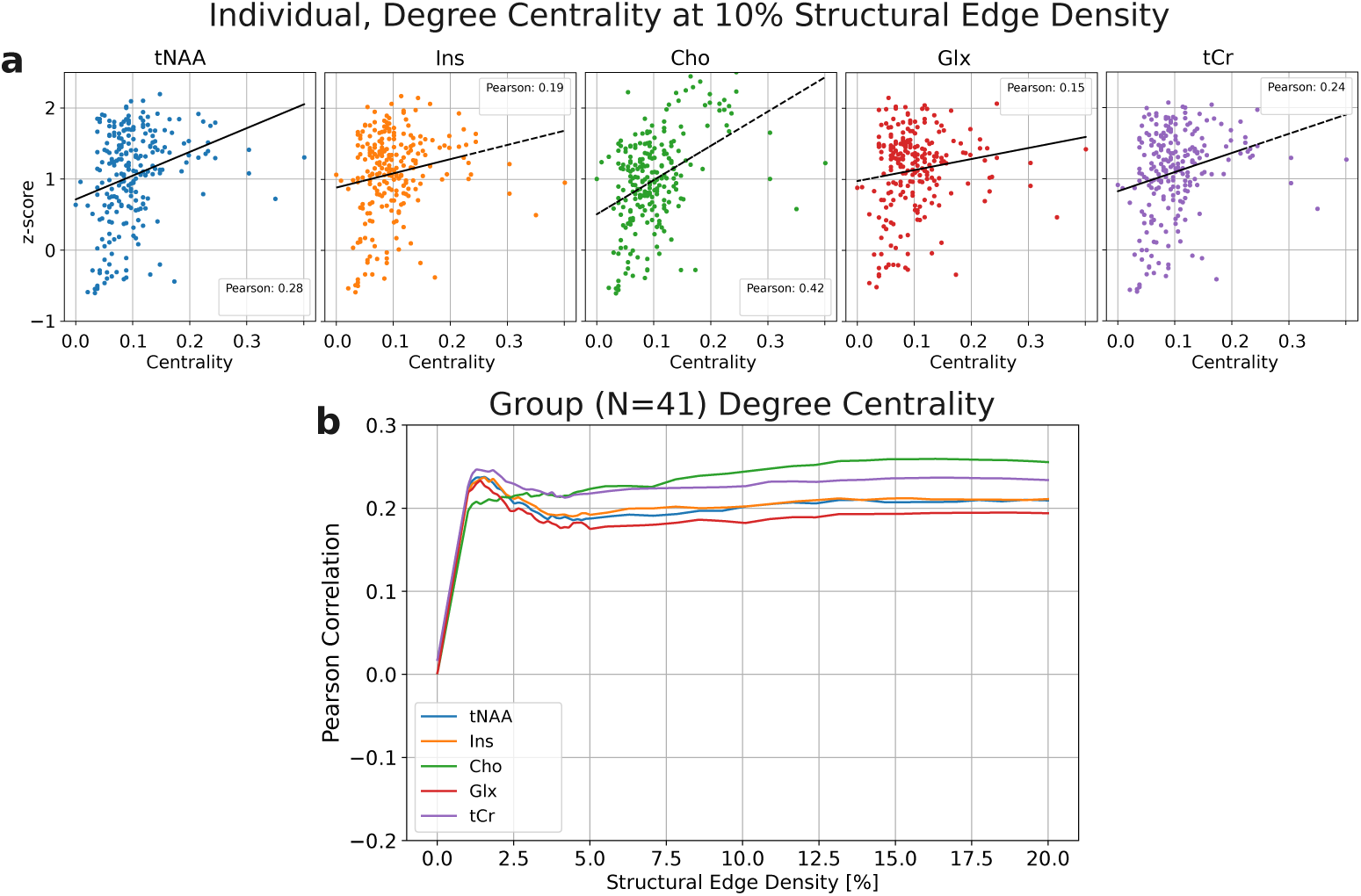
Metabolic Concentration and Structural Connectivity. **a**, Spearman correlation between five ^1^H-MRSI metabolite concentrations (z-score normalized) and structural node degree centrality at 10% edge connection density for an individual. **b**, Aggregated results of the Spearman correlation across 41 subjects between five ^1^H-MRSI metabolite concentrations (z-score normalized) and structural node degree centrality, evaluated at varying structural edge densities.

#### 2.4.2 Metabolic Similarity and Cytoarchitecture

To test the hypothesis that metabolic networks align with cytoarchitectonic similarity networks, we analyzed three datasets: the Cognitive-Consilience dataset, the Von Economo dataset, and the BigBrain dataset (see Methods 7.3.1).

The Cognitive-Consilience dataset, based on cortical laminar patterns Solari and Stoner (2011), was digitized by Seidlitz et al. Seidlitz et al. (2018) and extended in this study to include subcortical and cerebellar regions (see Methods 7.3.1), comprising nine bilaterally symmetric cytoarchitectonic classes (see Fig. 7a). We applied Gaussian Mixture clustering with nine clusters to the Geneva study group’s averaged MeSiM MSI map and compared the resulting clusters to the cytoarchitectonic classes. Alignment between the clusters and cytoarchitectonic classes was assessed using overlap metrics and spatial permutation tests (see Methods 7.4.5). Individual MeSiMs yielded a significant overlap score of 0.30 ± 0.02 (AdjPerm *p* = 0.008), and the group average MeSiM showed a higher score of 0.41 (AdjPerm *p <* 0.001). Cluster correspondence analysis revealed strong alignment in the motor cortex (class 1 and metabolic class 5, overlap: 0.47) and subcortical regions (class 8 and metabolic class 4, overlap: 0.91).

**Fig. 7.**
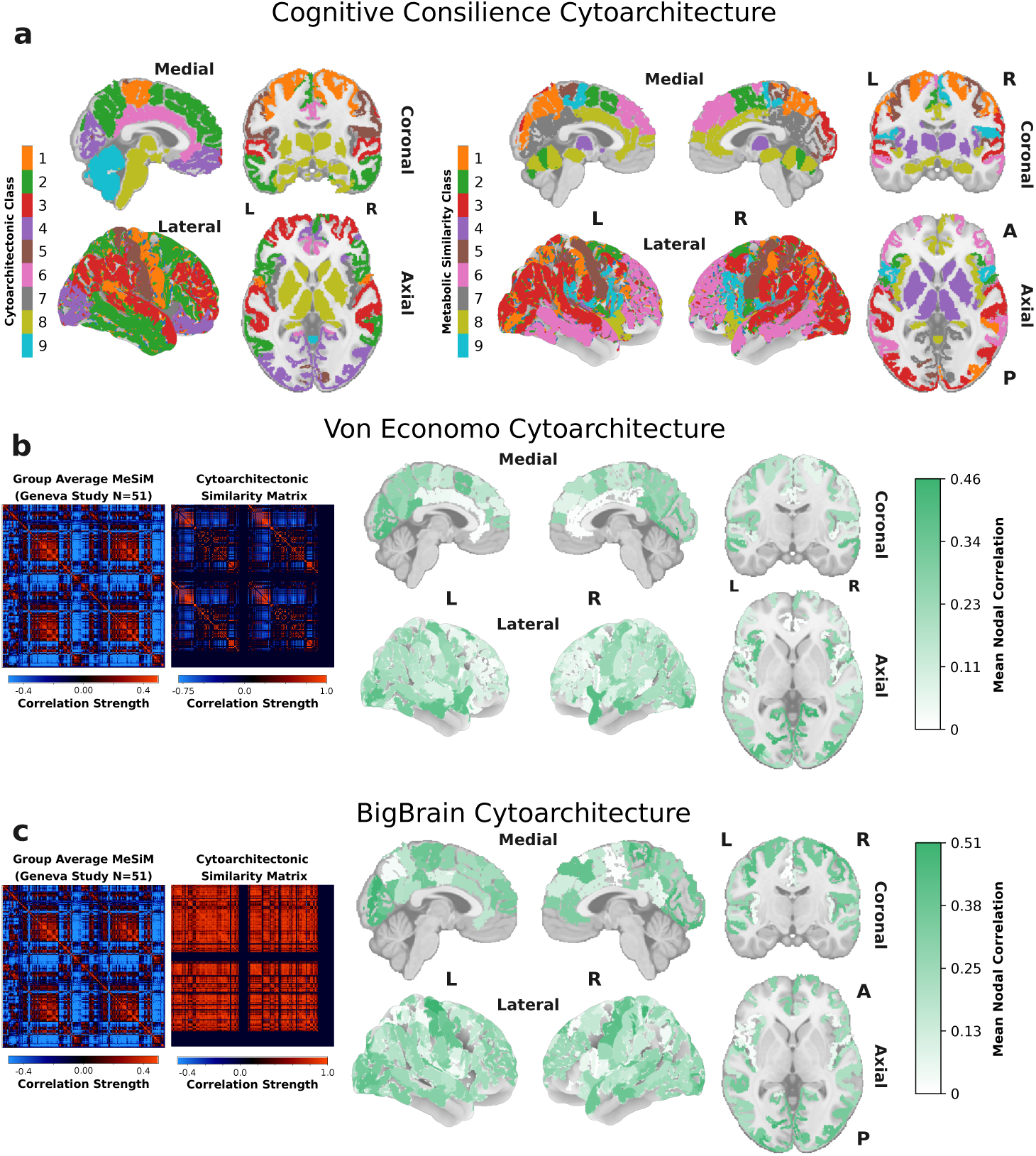
Metabolic Similarity and Cytoarchitectural Similarity. **a**, Cytoarchitectonic classification of cortical laminar patterns performed by Seidlitz et al. Seidlitz et al. (2018) (left) and based on the Cognitive-Consilience dataset Solari and Stoner (2011) and classification of metabolically similar cortical regions based . **b**, Von Economo cytoarchitectural similarity matrix and its corresponding absolute nodal correlation with the group MeSiM (*p <* 0.05 ; AdjPerm). **c**, BigBrain cytoarchitectural similarity matrix and its corresponding absolute nodal correlation with the group MeSiM (*p <* 0.05; AdjPerm).

The Von Economo dataset included 42 bilaterally symmetric cortical cytoarchitectonic profiles mapped onto the 210-node LFMIHIFIF-3 parcellation (left Fig. 7b). A direct edge-wise comparison between the group-average MeSiM and the Von Economo similarity matrix (see Methods 7.3.1) yielded a Spearman correlation of 0.10 (AdjPerm *p <* 0.001). This global association was primarily driven by the occipital lobe, medial superior prefrontal regions, precentral gyrus, superior temporal gyrus, and posterior insula (*r* = 0.45 − 0.51) as shown in the absolute nodal correlation metric on Fig. 7b (see Methods 7.5.7). In contrast, the anterior cingulate cortices showed minimal correlation.

The BigBrain similarity matrix (left Fig. 7c) Wei, Scholtens, Turk, and Van Den Heuvel (2018), derived from ultra-high-resolution cytoarchitectural data of the cortex Amunts et al. (2013) (see Methods 7.3.1), shows a node-wise correlation of 0.29 (AdjPerm *p <* 0.001) with the group-average MeSiM. The highest absolute nodal similarities were observed in the occipital lobe, postcentral gyrus, superior frontal gyrus (medial part), caudal-medial superior frontal cortex, and posterior cingulate cortex (*r* = 0.45 − 0.51). Similar to the Von Economo similarity matrix, the anterior cingulate cortex showed minimal correlation. These results suggest that MeSiM topology partly reflects spatial organizations of cytoarchitectonic patterns.

#### 2.4.3 Metabolic Similarity and Genetic Co-Expression

We next tested our second biological hypothesis, that MeSiM edges are characterized by high levels of gene co-expression, by leveraging the nearly complete human genome dataset (20,737 genes) from the Allen Brain Institute. Using data from six adult human post-mortem brains Hawrylycz et al. (2012), we mapped whole-genome transcriptional profiles onto the same LFMIHIFIF-3 parcellation used to define the MeSiM’s 277 nodes (see Methods 7.3.2). This approach allowed us to estimate inter-regional co-expression patterns (see Fig. 8) for every possible pair of nodes within the same anatomical reference frame as the MeSiM. A significant positive correlation was observed between MeSiM edge weights and whole-genome co-expression values (PermAdj *r* = 0.46, *p <* 0.001), indicating that metabolically similar regions also share similar gene expression profiles. Regions in the occipital lobe and superior frontal cortex (medial) showed the strongest absolute nodal correlation (*r* = 0.60 − 0.74), while the anterior cingulate cortex, brainstem and cerebellar regions contributed the least. After a gene ontology (GO) analysis of the 20,737 genes we extracted 250 GO categories with enriched contributions (FDR-corrected enrichment ratio *>*1). A leave-one-GO-class-out analysis, identified via the elbow method, 17 parent GOs that contributed the most (see Extended Data 8.6). These categories spanned three biological themes: cellular processes (e.g., cell division, ion transport), developmental processes (e.g., sensory organ development, neuron migration), and metabolic processes (e.g., membrane lipid biosynthesis). Categories related to immune regulation, structural processes, and DNA maintenance contributed the least. Collectively, these results indicate that MeSiM topology aligns with spatial expression patterns of brain-expressed genes that are essential for normal neuronal functions.

**Fig. 8.**
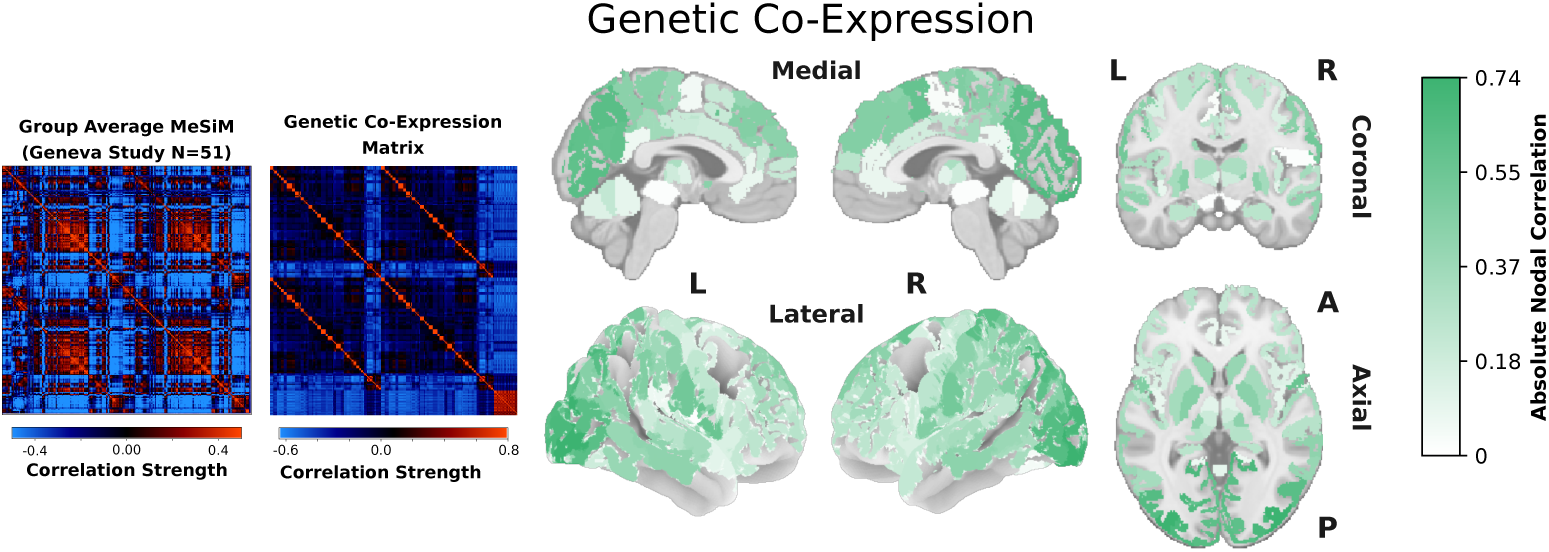
Metabolic Similarity and Genetic Co-expression. Genetic co-expression matrix (derived from Allen Human Brain atlas) and its corresponding absolute nodal correlation with the group MeSiM (*p <* 0.05 ; AdjPerm).

## 3 Discussion

We have demonstrated how MeSiMs, derived from 3D high resolution whole-brain metabolite imaging, can be used to estimate the metabolic similarity between brain regions in a sample of healthy participants at both the individual and group levels. We developed a reliable and consistent method for deriving MeSiMs, which we have shown to be also replicable in an independant sample scanned using a different MRI scanner. Furthermore, we outlined a novel brain metabolic network analysis, which enabled us to first determine the biological basis of the organization of metabolic networks derived from MeSiMs and test several hypotheses about their resemblance to other known networks.

Consistent with the complex topological organization observed in other naturally embedded networks Barabási (2013), MeSiMs reveal a modular network of nodes where spatially proximate nodes are metabolically similar, while distant nodes are metabolically dissimilar. Beyond this spatial tendency, we observed alignment along functional dimensions, with clustering of functionally similar regions (e.g., the limbic system) and segregation of functionally distinct regions (e.g., primary somatosensory vs. primary motor cortices). Integration across these distinct regions is mediated by metabolically high-degree connector hub nodes and whose spatial pattern aligns with a gradients of metabolic concentrations captured through the construction of metabolic fibers. Notably, this pattern is primarily driven by a smooth decrease in the tNAA/Cho ratio (relative to colinear levels of tCr, Glx, and Ins) from the occipital lobe (high ratio) through the parietal and frontal cortices, extending into subcortical structures, the brainstem, and the cerebellum (low ratio).

This decrease is consistent with existing MRS literature that demonstrated higher tNAA (a neuronal density marker) in gray matter–rich cortical regions but lower tNAA (and thus higher Cho, a membrane turnover marker) in white matter–rich subcortical areas such as the basal ganglia and brainstem Guan et al. (2017); Ratai et al. (2018); Zimny et al. (2013). The decrease in tNAA/Cho ratio from the occipital lobe to subcortical structures, reflects the shift from areas of higher neuronal density to those dominated by axonal content. Thus MeSiMs could potentially indicate the presence and extent of neurodegeneration, as a lower tNAA/Cho ratio reflects the loss of neuronal integrity and elevated membrane turnover in affected regions Guan et al. (2017); Ratai et al. (2018).

The pattern of MeSiM network organization follows a striking resemblance with the smoothly varying tonotopy along the basilar membrane of the cochlea, where the gradient of membrane properties (stiffness, width, and mass distribution) determines the spatial mapping of variable frequency responses of outer hair cells along the cochlear spiral von Békésy (1970). Here, the MSI gradient via the metabolic fiber reveals a smooth spatial variation in metabolic concentrations following the caudo-rostral axis and which determines the spatial mapping of distinct functional brain specializations. Thus, this ascertains and concludes the biological validity of the metabolic connectome as constructed from MRSI.

We observed that MeSiMs show poor correlation with structural connectivity, both directly (i.e., a structural edge does not imply metabolic similarity) and indirectly (i.e., intermediate connector nodes do not mediate correspondence), suggesting that the metabolic network represents an independent organizational framework distinct from structural connectivity pathways reliant on white matter tracts. As hypothesized, cortical areas connected in MeSiM networks exhibit a high degree of cytoarchitectural similarity and genetic co-expression. This pattern aligns with the established link between heightened co-expression and reliance on similar cellular machinery, which underlies parallel patterns of metabolic demand and activity Hansen et al. (2023); Oldham, Horvath, and Geschwind (2006); Wei et al. (2018).

In addition to illustrating a continuum of metabolic similarity across both proximal and distant brain regions, the metabolic fiber may carry deeper significance. Because it is constructed directly from the organizing pattern of the metabolic connectome and aligns closely with genetic co-expression patterns as well as cytoarchitectural organization, we propose that it is a remnant of the neurodevelopmental processes governing the maturation of the neural tube and its subsequent unfolding into subcortical and cortical structures. During early neurodevelopment, the neural tube differentiates into three primary divisions from rostral to caudal; the prosencephalon (forebrain), mesencephalon (midbrain), and rhombencephalon (hindbrain), preserving a sequential arrangement such that initially adjacent regions remain proximal in the mature central nervous system. While each region adopts a distinct gene expression profile that guides its development, function, and metabolism, the extent of its spatial variation remains constrained by proximity effect: adjacent areas share similar microenvironments Shin, Ming, and Song (2014), exhibit local diffusion of signaling molecules (morphogens) Goulas, Betzel, and Hilgetag (2019), and display aligned epigenetic modifications (e.g., DNA methylation patterns) Guo et al. (2011). Spatially assimilated regions share similar gene expression (genetic homophily), hence produce comparable metabolic enzymes and display analogous metabolic activity Oldham et al. (2006). Consequently, proximity in the neural tube translates into adjacency in the mature metabolic fiber, yielding its unbroken continuity observed from the forebrain through the midbrain to the hindbrain. Finally, our findings support the hypothesis that ^1^H-MRSI-measured metabolic profiles underpin the structural core of the human connectome, with high-centrality nodes showing elevated concentrations of metabolites. These results align with prior PET studies Scholtens, Schmidt, de Reus, and van den Heuvel (2014); Vaishnavi et al. (2010), which reported correlations between metabolic activity and structural centrality. Although the metabolites measured here with ^1^H-MRSI are only indirectly linked to glucose metabolism, they are closely associated with energy demands driven by glucose metabolism in regions of high cellular density. Also higher metabolite concentrations in high-centrality nodes likely reflect the cellular demands required to sustain these integrative hubs, with metabolites indicating neuronal integrity (tNAA), energy buffering and storage (tCr), neuronal activity (Glx), membrane turnover (Cho), and glial activity (Ins) Horská and Barker (2010).

The proposed ^1^H-MRSI–based metabolic connectome offers key advantages over positron emission tomography (PET) approaches reliant on radioactive tracers for glucose metabolism or neurotransmitter activity Ceccarini, Liu, Van Laere, Morris, and Sander (2020); Finnema et al. (2015); Wang, Yan, Xiao, Zuo, and Jiang (2020). When coupled with functional MRI, hybrid PET/MRI systems integrate both fMRI and PET-based ^18^F-fluorodeoxyglucose uptake for simultaneous metabolic and functional mapping Shan et al. (2022); Stiernman et al. (2021), but require lengthy scans (60–90 minutes), pose greater motion artifact risks, incur higher costs Pichler, Wehrl, Kolb, and Judenhofer (2008); Sander, Hansen, and Wey (2020), and involve neurovascular coupling effects Shan et al. (2022) and ionizing radiation—factors that limit their clinical practicality. Unlike invasive, single-process PET, MRSI is non-invasive, can quantify multiple metabolites in an acquisition time ranging from 8 to 20 minutes depending on the acquisition technique, and avoids ionizing radiation.

This study has several limitations to consider. MRSI’s limited coverage excludes orbitofrontal and basotemporal regions due to susceptibility artifacts and signal distortion. Additionally, the 5 mm isotropic spatial resolution challenges the accurate assessment of smaller subcortical regions. As a result, the MeSiMs constructed here may lack critical information, potentially omitting key network structures like the default mode network located in these areas. Furthermore, the constructed MSI map in this study was based on group-averaged data, which overlooks inter-individual variability. Factors such as participants’ resting or task states could influence metabolic connectivity and were not addressed here. Although we have not yet evaluated the alignment between MeSiMs and functional connectivity, our findings reveal that these networks possess a functional modular structure, which suggests that a strong correlation may exist. Our findings indicate that metabolic networks, through their associations with neurodevelopmental processes, may play a fundamental role in shaping both structural and functional brain systems. This highlights their potential as biomarkers, capable of revealing metabolic disruptions linked to disease even before structural or functional changes occur or symptoms manifest. Comparing these networks between controls and individuals at risk for disorders like schizophrenia could provide deep insights into underlying pathological mechanisms and identify metabolic alterations affecting central brain regions prior to the onset of the first psychotic episode. Thus, developing such networks should be viewed not merely as technical innovations but as practical tools poised for integration into a comprehensive suite of clinical applications.

## 4 Author Contributions

F.L. developed the methodology, performed the data analysis, conducted the experiments, contributed to data interpretation, and drafted the initial manuscript. E.C., P.S., and P.H. provided critical revisions. A.K. designed the MRSI sequence and provided critical revisions. Y.A-G. performed part of the data preprocessing and provided critical revisions. P.K. conceptualized the study, supervised the project, and provided critical revisions.

## 5#Acknowledgements

We gratefully acknowledge Jacob Seidliz for providing the digitized version of cytoarchitectonic classes based on the Cognitive-Consilience dataset, and Martijn van den Heuvel for providing the Big Brain cytoarchitectonic similarity matrix. Moreover, we gratefully acknowledge Jean-Baptiste Ledoux for having acquired all the MRI data from the Lausanne study, and Arnaud Merglen and Camille Marie Piguet for providing the MRI dataset of the Geneva study along with all their participants.

## 7 Methods

### Groups

#### Geneva Mindfulteen Study

Participants were recruited and scanned in the context of the Mindfulteen study, a randomized controlled trial to assess the effects of a mindfulness-based intervention on adolescent well-being. Details on recruitment, inclusion, and exclusion criteria can be found in the published protocol Piguet et al. (2022). Briefly, the study recruited adolescents (n = 69, 39 females, age range = 13-15 ± 0.8 yo) from the general population. Exclusion criteria included chronic somatic diseases or significant medical conditions, recent psychotherapy, recent use of psychotropic medications, and any DSM-IV psychiatric disorders, except for current anxiety disorders and/or a past episode of major depressive disorder. The study protocol was approved by the Geneva Regional Ethical Committee (CCER 2018-01731).

#### Lausanne Psychosis Cohort

For the replication of our results in an independent sample scanned at a different site, we included 13 healthy controls (4 females, ag range = 14-35 yo) from the Lausanne Psychosis Cohort Baumann et al. (2013). They were assessed by the Diagnostic Interview for Genetic Studies Preisig, Fenton, Matthey, Berney, and Ferrero (1999) to exclude a major mood, psychotic, or substance use disorder or had a first-degree relative with a psychotic disorder.

### 7.1 MRI Image Acquisition

For the Mindfulteen study MR data have been acquired on 3-Tesla scanner (Magnetom TrioTim, Siemens Healthineers, Forchheim, Germany) equipped with a 32-channel head coil at the Brain and Behavior laboratory in Geneva. Each scanning session includes a magnetization-prepared rapid acquisition gradient echo (MPRAGE) T1-weighted sequence with 1 mm in-planeresolution and 1.2 mm slice thickness, covering 240 × 257 × 160 voxels. The repetition (TR), echo (TE), and inversion (TI) times were 2300, 2.98, and 900 ms, respectively. The diffusion spectrum imaging (DSI) sequence included 128 diffusion-weighted images with a maximum b-value of 8000 s / mm^2^ and one b_0_ reference image. The acquisition volume was made of 96 × 96 × 34 voxels with 2.2 × 2.2 × 3 mm resolution. TR and TE are 6800 and 144 ms, respectively.

For the Mindfulteen study, magnetic resonance (MR) data were acquired using a 3-Tesla scanner (Magnetom TrioTim, Siemens Healthineers, Forchheim, Germany) equipped with a 32-channel head coil at the Brain and Behavior Laboratory in Geneva. Each scanning session included a magnetization-prepared rapid acquisition gradient echo (MPRAGE) T1-weighted sequence with an in-plane resolution of 1 mm and a slice thickness of 1.2 mm, covering a volume of 240 × 257 × 160 voxels. The repetition time (TR), echo time (TE), and inversion time (TI) were set to 2300 ms, 2.98 ms, and 900 ms, respectively. Additionally, a diffusion spectrum imaging (DSI) sequence was performed, acquiring 128 diffusion-weighted images with a maximum b-value of 8000 s/mm^2^ and one b_0_ reference image. The acquisition volume for DSI consisted of 96 × 96 × 34 voxels with a resolution of 2.2 × 2.2 × mm. The TR and TE for the DSI sequence were 6800 ms and 144 ms, respectively.

The 3D ^1^H-FID-MRSI sequence accelerated by compressed-sensing Klauser et al. (2021) was acquired with 1.50 ms TE, 372 ms TR and 35 deg flip angle. The Field-of-View (FoV) size was 210 × 160 × 105 mm (anterior-posterior, right-left, head-foot directions) with a 95mm-thick slab selection, with a spatial resolution of 5×5 × 5.3 mm and the spectral bandwidth was 2 kHz acquired with 512 points. The reference water acquisition had the following parameters: same TE, TR of 25 ms, lower flip angle of 3°, same FOV size, lower resolution of 6.6 x 6.7 x 6.6 mm, same bandwidth and free induction decay (FID) size of 16 points.

For the Lausanne Psychosis Cohort, MRI sessions were conducted on a 3-Tesla scanner (MAGNETOM Prisma fit, Siemens Healthineers, Forchheim, Germany) at the Lausanne University Hospital for 13 healthy controls. While the MPRAGE and DSI sequences parameters were identical to those used in the Mindfulteen study, the the 3D ^1^H-FID-MRSI sequence had different parameters than the one implented for the Geneva study (TE = 1.5 ms, TR = 372 ms and flip angle of 40 deg. The FoV was the same as in Lausanne, with the same spatial resolution of 5×5 × 5.3 mm, as the spectral bandwith and FID size. The water acquisition had the following parameters: same TE of 1.5 ms, TR of 36 ms, and a lower flip angle of 5 deg. FOV size, resolution, bandwidth and vector size were the same as for the Geneva study.

### 7.2 Preprocessing

#### 7.2.1 Anatomical

For each subject, the anatomical T1-weighted image was used to parcellate the brain into different regions using the FreeSurfer package (version 7.2.0, http://surfer.nmr.mgh.harvard.edu) and subsequently identify nine distinct supra-regions: cortex, basal ganglia, thalamus, amygdala, hippocampus, hypothalamus, cerebellum, brainstem, and white matter.

We relied on the Chimera parcellation software tool Alemán-Gómez (2024) to create a combined volumetric parcellation for each input subject anatomical image based on the following atlases (as listed in the original Chimera repository) and referred to as the LFMIHIFIF-3 parcellation scheme:

- Lausanne cortical parcellation scale 3 Cammoun et al. (2012),
- Basal ganglia parcellation Fischl et al. (2002),
- Thalamus parcellation Najdenovska et al. (2018),
- Amygdala parcellation Saygin et al. (2017)
- Hippocampus parcellation Iglesias et al. (2015)
- Hypothalamus parcellation Billot et al. (2020),
- Cerebellum parcellation Fischl et al. (2002),
- Brainstem parcellation Iglesias et al. (2015).

#### 7.2.2 DSI and Structural Connectivity

The DSI volumes were visually inspected for signal loss, which could indicate motion artifacts and potential exclusion from the analysis Yendiki, Koldewyn, Kakunoori, Kanwisher, and Fischl (2014). The DSI data were reconstructed following the methodology described by Wedeen et al. (2005) Wedeen, Hagmann, Tseng, Reese, and Weisskoff (2005). Preprocessing, anatomically constrained tractogram construction using deterministic tractography, and structural connectivity computation were performed using the MRtrix3 software tool Tournier et al. (2019).

Particularly, the peaks of the local fiber orientation distribution functions (fODFs) (maximum of three peaks per fODF) were computed and served as input for the deterministic tractography Tournier, Calamante, and Connelly (2012). The tractography parameters included a maximum track length of 300 mm, 32 seeds per white matter voxel per diffusion direction, and termination criteria of a 60-degree angle between subsequent propagation steps or reaching the white-gray matter boundary. To ensure reproducibility, 10^6^ streamlines were selected Newlin, Rheault, Schilling, and Landman (2023).

A weighted, undirected brain connectivity matrix was reconstructed for each subject by filtering the whole-brain tractogram using Spherical-deconvolution Informed Filtering of Tractograms (SIFT) Smith, Tournier, Calamante, and Connelly (2013), which refined streamline contributions based on their alignment with underlying diffusion data and termination coordinates. We defined the sub-regions of the Chimera parcellation (see Section 7.2.1) as the nodes of the structural connectivity. The strength of the connections *S_ij_* between any two node *i* and *j* were computed as the number of streamlines *ns_ij_* connecting these two regions weighted by the inverse of their average length 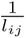 and the inverse of the node’s volumes 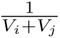 i.e. 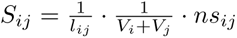 The length and volume terms were used to eliminate the bias towards longer fibers introduced by the tractography algorithm and the bias for the variable size of cortical ROIs respectively Hagmann et al. (2008). For consistency and to limit possible false-positive connections, the connections that present in less than 50% of participants were discardedde Reus and van den Heuvel (2013); Zalesky et al. (2016).

#### 7.2.3 MRSI

The MRSI data was reconstructed using a low-rank model constrained by total-generalized variation, with prior removal of subcutaneous lipid contamination and residual water signals Klauser et al. (2021). Following reconstruction, the spatio-spectral data was analyzed using LCModel Provencher (2001) to quantify metabolite signal in each voxel, using the water signal from the additional acquisition as a reference.

The basis set for LCModel fitting included the following metabolites: N-acetylaspartate (NAA), N-acetylaspartylglutamate (NAAG), creatine (Cr), phospho-creatine (PCr), glycerophosphocholine (GPC), phosphocholine (PCh), myo-inositol (mI), scyllo-inositol (sI), glutamate (Glu), glutamine (Gln), lactate (Lac), gamma-aminobutyric acid (GABA), glutathione (GSH), taurine (Tau), aspartate (Asp), and alanine (Ala). Only a subset of these metabolites could be reliably resolved and due to overlapping spectral peaks, certain metabolites were combined: NAA and NAAG (denoted tNAA), Cr and PCr (denoted tCr), GPC and PCh (denoted Cho), mI (denoted Ins), and Glu and Gln (denoted Glx), resulting in five distinct metabolite volumes. LCModel also provided spectral quality metrics, such as the signal-to-noise ratio (SNR), Cramer-Rao Lower Bound (CRLB) for each metabolite estimation.

#### 7.2.4 MRSI Volume Parcellation

The MRSI tCr volume was co-registered to the T1-weighted image using Advanced Normalization Tools (ANTs) Tustison et al. (2021), where mutual information served as the cost function for optimization. This resulted in a combination of an affine transform and a symmetric image normalization transform that was used to project the Chimera parcellation into the MRSI subject space. This procedure was repeated for each subject’s ^1^H-MRSI recording, resulting in a personalized MRSI parcellation and yielding five MRSI volumes (tNAA, tCr, Cho, Ins, and Glx) distributed across 277 brain regions.

The multi-scale cortical parcellation proposed by Cammoun et al. Cammoun et al. (2012) was selected for its hierarchical organization, which allows for flexible regional granularity. This parcellation provides coarse-scale segmentations with fewer but heterogeneously sized regions (scale 1) as well as finer-scale subdivisions with a greater number of parcels of relatively uniform size (scale 5). Due to the limited spatial resolution of ^1^H-MRSI data (5 mm isotropic) compared to the higher resolution of T1-weighted anatomical images (1 mm), scale 3 was chosen. This scale offers a more uniform distribution of parcel sizes, which better matches the resolution of the MRSI data while preserving anatomical specificity.

#### 7.2.5 Estimation of Metabolic Profiles

After parcellation, regions smaller than 8 MRSI voxels (<1 cm^3^) or those with poor MRSI coverage in more than 70% of subjects were excluded from the 277 regions, resulting in *N* = 210 retained regions. The discarded regions were mainly located in orbitofrontal and baso-temporal regions where ^1^H-MRSI has limited coverage due to susceptibility artifacts. These factors reduce signal-to-noise ratio and introduce spectral contamination, hindering reliable metabolite quantification in these areas. Additionally, the ^1^H-MRSI sequence covered only the superior portion of the cerebellum. The complete ^1^H-MRSI mask coverage is shown in the Extended Data 8.1. Median metabolite values were calculated inside a given parcel and z-score normalized across regions, yielding a 5-feature metabolic profile per region. To address suboptimal spectral peak resolution, we repeated the process *K*_pert_ = 50 times to generate perturbed versions of the original metabolic profile. Specifically, for each voxel (*i, j, k*), the metabolite concentration was resampled from a normal distribution with mean equal to the original MRSI signal *Y_ijk_* and standard deviation equal to 3*σ_ijk_*^2^, where *σ_ijk_*^2^ is the Cramér-Rao Lower Bound (see Methods 7.2.3) derived from the LCModel. Mathematically,

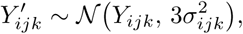

where *Y_ijk_* is the original MRSI signal value. This augmentation increased the sample size of the metabolic profiles by a factor of *K*_pert_, providing *K*_pert_ − 1 additional perturbed profiles for each brain region, in addition to the original profile.

#### 7.2.6 Metabolic Similarity Index

To visualize the spatial embedding of the 210×210 dimensional group-averaged MeSiM (Geneva sample, *N* = 51), we reduced each of its 210 high-dimensional feature vectors **x***_i_*—corresponding to the parcel connectivity profiles—into a lower-dimensional representation. First, Principal Component Analysis (PCA) was applied for denoising and variance capture, resulting in a 50-dimensional embedding. Subsequently, t-distributed Stochastic Neighbor Embedding (t-SNE) was employed to further reduce the dimensionality to obtain the metabolic similarity inidex (MSI) *µ_i_* at every nodes *i* . As prescribed by Kobak and Berens (2019), PCA initialization was used to stabilize and improve the convergence of t-SNE, which in turn was chosen to preserve the observed local connectivity patterns of the MeSiM.

t-SNE converts high-dimensional distances between data points into conditional probabilities (or similarities) and then maps them into a low-dimensional space. A crucial parameter in t-SNE is perplexity, which can be interpreted as a measure of how many close neighbors each point has. It also balances local and global connectivity patterns, where lower perplexity emphasizes local structures at the expense of global structure, and higher perplexity does the opposite. The method adjusts the lowdimensional output features *µ_i_* by minimizing the Kullback–Leibler (KL) divergence between the probability distributions in the high-dimensional and low-dimensional spaces, as computed via a t-Student-based metric.

We explored perplexity values from 20 to 60 and observed minimal changes in the KL divergence (ranging approximately from 0.55 to 0.42). Consequently, following Kobak and Berens (2019), we chose a perplexity of 30 to strike a balance between preserving both local and global structures. Finally, an early exaggeration factor of 12, as suggested by Linderman and Steinerberger (2019), was applied to enhance cluster separation during the initial stages of optimization. From these parameters, we reconstructed a filtered version of the original MeSiM, whose elements *q_ij_* were computed using the t-Student-based metric:

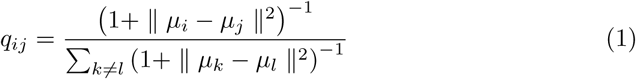

where *µ_i_* is the dimension-reduced representation of the *i*-th node’s connectivity profile *x_i_*. We then rescaled these values to 2*q_ij_* − 1 to restore the original MeSiM value range [−1, 1] and verify the overall pattern conservation of the original MeSiM shown in Supplementary Fig. 2.

#### 7.2.7 Metabolic Fibre Construction

In diffusion MRI tractography white matter streamlines are traced by aligning their tangent vector to the principal eigenvectors field of the diffusion tensor field, weighted by regions of strong fractional anisotropy and thereby structurally connecting distant gray matter nodes. Here we define metabolic paths by aligning their tangent vector with the MSI gradient weighted by regions of high metabolic density and consequently *relaying* metabolically distinct regions.

Instead of identifying a continuous curve in space, the path we aim to construct is a sequence of edges in a discrete lattice (finite set of nodes) within the metabolic network. We proceed to introduce the definitions for a path, the MSI gradient, and nodal metabolic strength.

Let *G* = (*V, E, w*) be a weighted network where *V* is a finite set of nodes, *E* ⊆ *V* ×*V* is a set of edges and *w* : *E* 1→ [−1, 1] is the weight function which assigns a MeSiM value *w_ij_* to nodes *i* and *j*. We define a path *γ* of length *L* as an ordered sequence of nodes

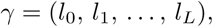

such that for each integer *t* (0 ≤ *t < L*), the consecutive pair (*l_t_, l_t_*_+1_) forms an edge in the network. We designate *l*_0_ as the start node and *l_L_* as the end node. We define the MSI gradient between node *i* and *j* as the absolute spatial variation of the their respective MSI values:

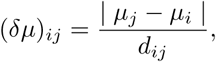

where *µ_i_* is the MSI value at node *i*, and *d_ij_* = ‖**r***_j_* − **r***_i_*‖ is the Euclidean distance between nodes *i* and *j*. The nodal (metabolic) strength of node *i* is defined as the sum of its adjacent connections:

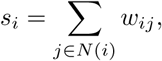

where *N* (*i*) is the set of adjacent gray matter nodes of node *i* and where adacency here does not relate to metabolic adjacency (high *w_ij_*) but rather to physical adjacency i.e. a parcel *j* is considered adjacent if it contains at least one voxel that directly neighbors a voxel in parcel *i*. We require *γ* to follow the arrangement of potential nodes that are both:

- **Condition 1** highly metabolically similar to their proximal neighbors (i.e. minimize MSI gradient)
- **Condition 2** highly connected (i.e., maximize nodal strength).

The first condition is analogous to optimally aligning the tangent vector of a smooth curve in continuous space with a vector field (MSI gradient), while the second condition prioritizes highly connected nodes so that the path captures the principal variability of the network. We formulate the edge-based cost function *c*(*i, j*)

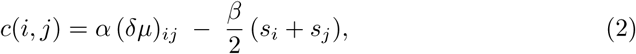

where *α >* 0 and *β >* 0 are trade-off parameters controlling the relative importance of minimizing the MSI gradient versus favoring highly connected nodes. Then, for a given path *γ* = (*l*_0_*, . . . , l_L_*), its mean total cost is

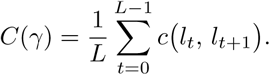

where *γ*^∗^ minimizes:

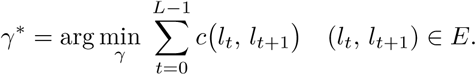

We employed a brute-force algorithm via Depth-First Search backtracking to:

1. identify all possible paths connecting the start and end nodes,
2. calculate the cost *C*(*γ*) induced by each path,
3. select the paths which yielded the minimal cost value.

Given the hemispheric symmetry of the MSI map, pathfinding was performed independently in each hemisphere. In both cases, the precuneus (PCN; lowest index) served as the starting node, and the rostral anterior cingulate cortex (rACC; highest index) as the ending node. Because subcortical regions form a separate gray matter network, we applied the same pathfinding approach to these regions by splitting each hemisphere while keeping the brainstem as a shared structure.

Figure Supplementary Fig. 4 displays the total cost function at each node iteration *t* for *α* = 1 and *β* ∈ [0, 1]. Here, we identified 32,865 and 51,709 possible paths in the left and right hemispheres of the neocortex, respectively, and 2,463 and 1,590 paths in the left and right hemispheres of the subcortical regions. From these, we selected the paths yielding the minimal total cost (colored curves). In the neocortex of both hemispheres, this procedure resulted in two main paths (the lateral parietal path and the superior parietal path), as well as one subcortical path connecting the cerebellum and the caudate nucleus via the brainstem’s pontine region.

### 7.3 Datasets

#### 7.3.1 Cytoarchitecture

#### Congitive Consilience Classification

We acquired three distinct cytoarchitectonic datasets, referred to as the cortical-laminar, Von Economo, and BigBrain cytoarchitectures. The first cytoarchitecture is based on an independent modular decomposition representing the five classic cortical laminar patterns described by Von Economo and Koskinas von Economo and Koskinas (1925). Following the approach of Seidlitz et al. Seidlitz et al. (2018), we manually assigned each node from our 308-region parcellation to one of these five cortical classes, guided by the methodology outlined by Solari and Stoner Solari and Stoner (2011). The cingulate and insular cortices were assigned respectively to a sixth and a seventh class. In addition, we categorized subcortical regions—including the thalamus, hypothalamus, amygdala, hippocampus, and brainstem—into a distinct eighth class to account for their unique cytoarchitectonic characteristics. Finally, the cerebellum parcels were assigned to a ninth class, as they are composed of distinctive cell types (e.g., Purkinje cells) found exclusively in this region of the brain.

#### Von Economo Similarity Matrix

The Von Economo cytoarchitecture van den Heuvel, Scholtens, Barrett, Hilgetag, and de Reus (2015) represents a digitized version of the original cytoarchitectonic brain profiles described by Von Economo and Koskinas von Economo and Koskinas (1925), forming a comprehensive brain atlas. This atlas comprises 48 ‘most important‘ distinct cortical areas and provides detailed layer-specific histological information, including neuronal count, neuron size, and cortical thickness. The Von Economo cytoarchitectonic similarity matrix was constructed as a pairwise Pearson correlation matrix of all 48 profiles.

#### Big Brain Similarity Matrix

The BigBrain cytoarchitectonic profiles Wei, Scholtens, Turk, and Van den Heuvel (2019) were derived from the ultrahigh-resolution BigBrain dataset Amunts et al. (2013) (see Methods 7.3.1), where cortical profiles capturing the laminar cell number and density of the cortex were extracted. The resulting BigBrain similarity matrix, including pairwise similarity scores between *N* brain regions (excluding subcortical and cerebellar regions), was directly shared with us by the authors of Wei et al. Wei et al. (2019).

#### 7.3.2 Genetic Co-Expression

Gene expression data were obtained from the publicly available dataset provided by the Allen Institute for Brain Sciences (AIBS). To reduce redundancy in complementary RNA (cRNA) hybridization probes measuring overlapping gene expression, expression values were averaged across probes targeting the same gene, while probes without matched genes were excluded. This process resulted in a dataset comprising 20,152 genes across 3,702 samples.

Given the symmetry in gene expression observed between hemispheres Pletikos et al. (2014), AIBS data include samples from both hemispheres for only two subjects. However, due to the under-sampling of the right hemisphere, all analyses were restricted to the left hemisphere (n = 152 regions). Gene expression data for each subject were mapped onto the fsaverage (MNI152) volumetric template. This was achieved by assigning samples to the nearest centroid within the left hemisphere regions using T1-weighted MRI scans from each AIBS subject. For the two subjects with right hemisphere data, coordinates of the right hemisphere samples were reflected before mapping to ensure consistency.

Median gene expression values were calculated for each region across participants (*N* = 6) and standardized using z-scoring, resulting in a 152 × 20,152 matrix representing genome-wide expression across 152 left hemisphere regions. A 152 × 152 gene co-expression matrix was then generated by computing pairwise Pearson correlations of gene expression between each region. This co-expression matrix, along with regional expression values, was used for comparisons with the left hemisphere data of the group MeSiM. Gene ontology enrichment analysis was performed using GOATools Klopfenstein et al. (n.d.)

### 7.4 Statistical Analysis

#### 7.4.1 MeSiM Variablity

The variability of a set of metabolic similarity matrices was assessed by calculating the mean and standard deviation of variances across the set, and then performs a t-test to determine if the observed dispersion is significantly different from zero.

The variability of a set of metabolic similarity matrices with respect to the group average was determined by calculating the Pearson correlation coefficients between each matrix and the group average matrix. The resulting p-values were adjusted for multiple comparisons using the Benjamini-Hochberg method, yielding the average and standard deviation of the Fisher Z-transformed correlations, along with an overall p-value.

#### 7.4.2 Adjacent Parcel Permutation

To assess statistical significance when comparing spatially embedded datasets (e.g., similarity matrices, brain region classification scores) while preserving local spatial properties, we employed a customized spatial permutation method termed AdjPerm. This approach maintains local topology by restricting permutations to occur only between adjacent regions. Specifically, we defined adjacency based on anatomical proximity, considering each brain parcel to be adjacent to its immediate neighboring parcels. During each permutation, data values were randomly shuffled among these adjacent parcels, preserving local spatial dependencies. We fixed the number of unique permutations to 1000. By maintaining local topology, this method effectively accounted for inherent spatial autocorrelation without introducing anatomically implausible configurations.

#### 7.4.3 Volumetric Spin Test

To assess statistical significance when comparing spatially embedded datasets (e.g., similarity matrices, brain region classification scores) while preserving global spatial properties, we employed a customized spatial permutation method, termed VolSpin, analogous to the spin test Alexander-Bloch et al. (2018), adapted for volumetric data. We generated random rotation matrices to rotate the Euclidean coordinates of the centroids of each brain parcel along two concentric volumetric spheres: the first containing cortical regions and the second containing subcortical regions. This approach ensured that cortical parcels were always mapped to other cortical parcels and subcortical parcels to subcortical ones, thereby preserving the brain’s global topology. Additionally, we maintained hemispheric consistency by rotating parcels within each cerebral hemisphere so that they remained in the same hemisphere post-rotation. After rotation, we reassigned data values by mapping the rotated coordinates back to the original brain parcels, matching each to the parcel with the closest Euclidean distance. To avoid introducing anatomically implausible configurations and creating replicate samples in the null distribution, we verified the uniqueness of each rotated configuration and discarded any duplicates. We fixed the number of unique permutations to 1000.

#### 7.4.4 Normalized Mutual Information Score

We evaluated the agreement between two 3D scalar maps using the Normalized Mutual Information score. If both datasets were spatially embedded, we supplemented the significance of the overlap score with the AdjPerm test (see Methods 7.4.2).

#### 7.4.5 Inter-Class Overlap Score

We evaluated the agreement between two labeled datasets using the Hungarian algorithm (Kuhn-Munkres algorithm). This approach identifies an optimal one-to-one mapping between the sets of labels, maximizing the number of samples that are consistently classified under this mapping. We supplemented the significance of the overlap score with the AdjPerm test (see Methods 7.4.2).

### 7.5 Network Based Statistics

#### 7.5.1 Weighted Matrix Binarization

Network graphs were generated by binarizing weighted matrices (similarity or connectivity) based on a specified edge density *ρ*. For matrices containing negative values, negative weights were rectified to their absolute values before applying the binarization threshold. This threshold was chosen to achieve the desired proportion of binarized connections *ρ*. Graph analyses were conducted across a wide range of edge densities (1 to 30 %) for each weighted matrix.

#### 7.5.2 Rich-Club

We computed the rich-club coefficient *ϕ*(*k*) for a given binarized network as

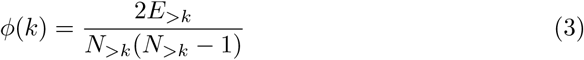

where *E_>k_* is the number of edges among nodes with a degree greater than *k*, and *N_>k_* is the number of such nodes.

#### 7.5.3 Degree Centrality

We calculated the centrality of a node using **degree centrality**, which measures the number of direct connections a node has to other nodes in the network. For a node *i*, the degree centrality *C_D_*(*i*) is defined as:

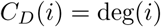

where deg(*i*) represents the number of edges connected to *i*. Degree centrality highlights the immediate influence of a node based on its local connectivity.

#### 7.5.4 Random Networks

For each binarized graph, we assessed the statistical significance of the observed network metrics by generating an ensemble of 1,000 randomized networks using the Maslov-Sneppen rewiring procedure, which preserves the number of nodes, total number of edges, and the degree distribution of the original graph while randomizing the specific wiring of edges. We then computed empirical p-values for each topological metric by comparing the observed value to the distribution of corresponding values across the 1,000 random graphs. Specifically, for each metric, the p-value was calculated as the fraction of randomized networks in which the metric was greater than or equal to the observed value.

#### 7.5.5 Higher Order Connectivity

The higher-order connectivity of a weighted or binarized network matrix **A** was computed as the *k*-th order cosine similarity of **A** with itself:

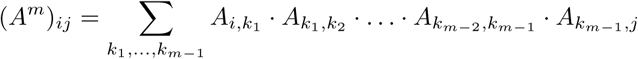

where Σ*_l,m_ A_k,l_* · *A_m,n_* denotes the cosine product of the *l*-th node feature with the *m*-th node feature of **A**.

#### 7.5.6 Network Communicability Model

To compute the influence of distant nodes from a structural connectivity matrix **S** on the state of the MeSiM **M**, we relied on a communicabilty model Kondor and Lafferty (2002) and estimated the Green’s function of the network heat equation , which corresponds to the communicability expression:

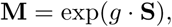

The decay parameter *g* controls the influence of distant nodes structural edges on the metabolic connectivity and was estimated by minimizing the mean squared loss between the model’s output (**M**) and the observed metabolic matrix (**M**_obs_). Predictive performance was then quantified as the mean Spearman correlation between the predicted MeSiM (**M**) and the observed MeSiM (**M**_obs_) given the optimal *g*.

#### 7.5.7 Absolute Nodal Correlation

When comparing the nodal similarity between two matrices **A** and **B** of *N* nodes, we define the absolute nodal correlation of node *i* as the absolute value of the Pearson correlation between its connectivity profiles in **A** and **B**. Here, the connectivity profile of a matrix **X** is given by

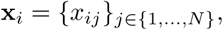

which is the row vector containing all similarity measures of node *i* with the other nodes in the network.

## 8 Extended Data

### 8.1 MeSiM Connectome Coverage

After individual ^1^H-MRSI volume parcellations, regions with fewer than 10 ^1^H-MRSI voxels in more than 70% of subjects were excluded, resulting in the effective MeSiM coverage shown in Supplementary Fig. 1. The excluded regions were primarily located in the orbitofrontal and basotemporal areas, where ^1^H-MRSI coverage is limited due to susceptibility artifacts and signal distortion. Additionally, the inferior part of the cerebellum was excluded due to the spatial cutoff of the ^1^H-MRSI sequence.

### 8.2 Reconstructed MSI Matrix

### 8.3 Impact of Cortical Parcellation Schemes on MeSiM Construction

We evaluated the impact of different cortical parcellations, specifically Schaefer200 Schaefer et al. (2018) (200 cortical parcels) and MIST197 Urchs et al. (2019)(197 cortical parcels), on the construction of MeSiMs. These parcellations, derived from functional cortical atlases, were chosen to contrast the structurally based Lausanne parcellation in the Chimera LFMIHIFIF-3 scheme, given their comparable number of cortical parcels. Both were integrated into the Chimera schema by substituting the Lausanne parcellation with Schaefer-200 and MIST197. The Gini coefficient, commonly used to assess inequality, was applied here as a measure of uniformity in parcel sizes. Lower Gini values indicate more uniformly sized parcels. The Schaefer-200 (0.21), MIST197 (0.22), and Lausanne (0.20) parcellations all exhibited low and comparable Gini coefficients, demonstrating their suitability for schemes requiring uniformly sized parcels. Because different parcellation schemes lead to differently shaped and differently mapped matrices, they cannot be directly compared. Hence, we directly compared their resulting MSI maps, with the original MSI map that resulted from the LFMIHIFIF-3 parcellation scheme using the normalized mutual information (NMI), and supplemented the observed score with PermAdj. After discarding parcels with poor ^1^H-MRSI coverage, Schaefer-200 and MIST197 retained 188 and 177 parcels, respectively, compared to 192 in the LFMIHIFIF-3 atlas. Both alternative atlases showed high NMI with LFMIHIFIF-3 (0.80 for MIST197, 0.85 for Schaefer-200; both *p <* 0.05 PermAdj), indicating their equivalence in generating MSI maps.

### 8.4 Rich-Club Analysis

Rich-club coefficients were computed for the MeSiM matrices of all individuals in the Geneva sample and tested against 1,000 Erdős-Rényi random graphs across edge densities ranging from 1% to 30% (see Supplementary Fig. 3a). The same analysis was applied to the group-averaged MeSiM (*N* = 51) (see Supplementary Fig. 3b). Rich-club coefficients were significant across all subjects and the averaged MeSiM.

### 8.5 Metabolic Fibre Cost Function

### 8.6 Gene Enrichment Cutoff

**Supplementary Fig. 1.**
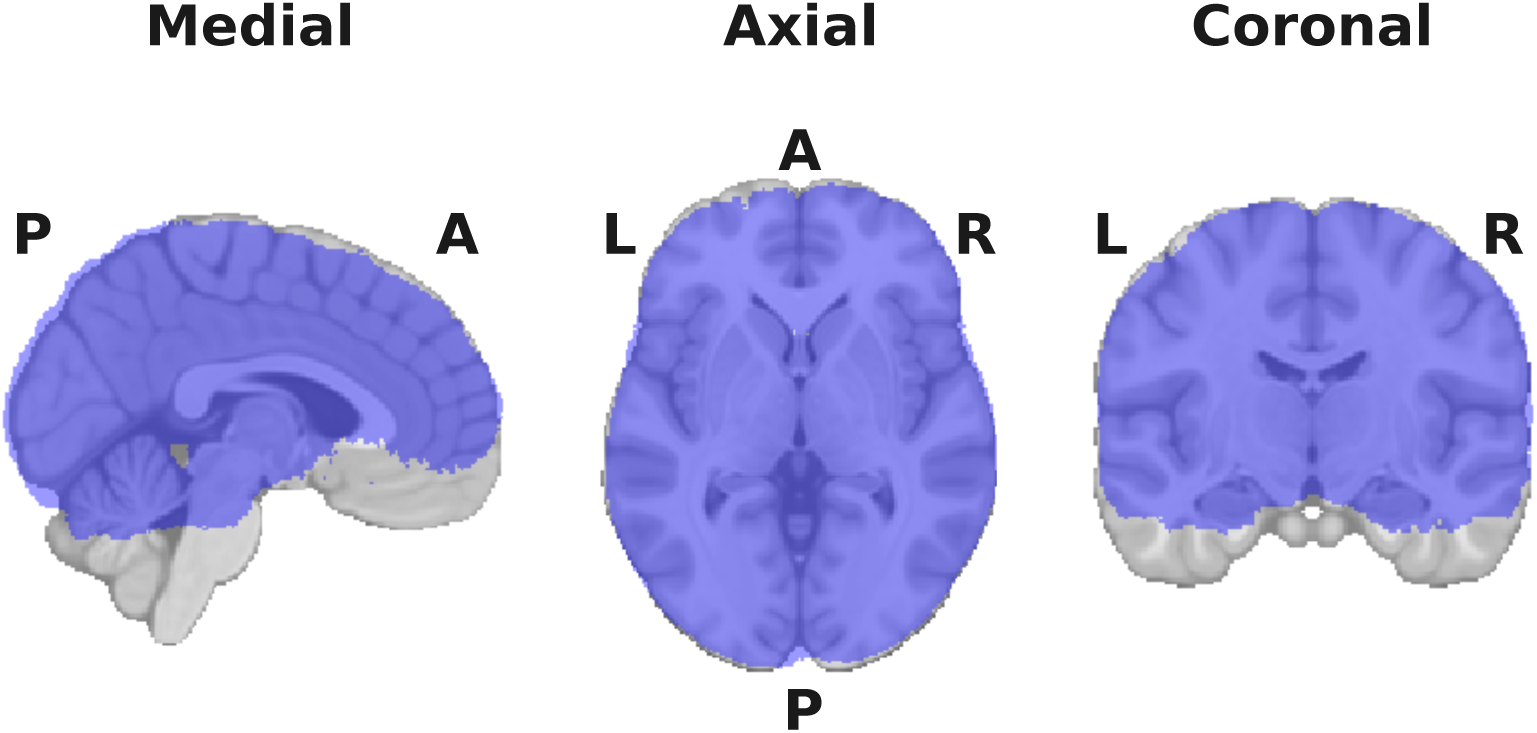
MeSiM Coverage. Effective MeSiM coverage for the Geneva sample (*N* = 51), excluding regions with limited ^1^H-MRSI voxels (orbitofrontal, basotemporal) and the inferior cerebellum due to sequence cutoff.

**Supplementary Fig. 2.**
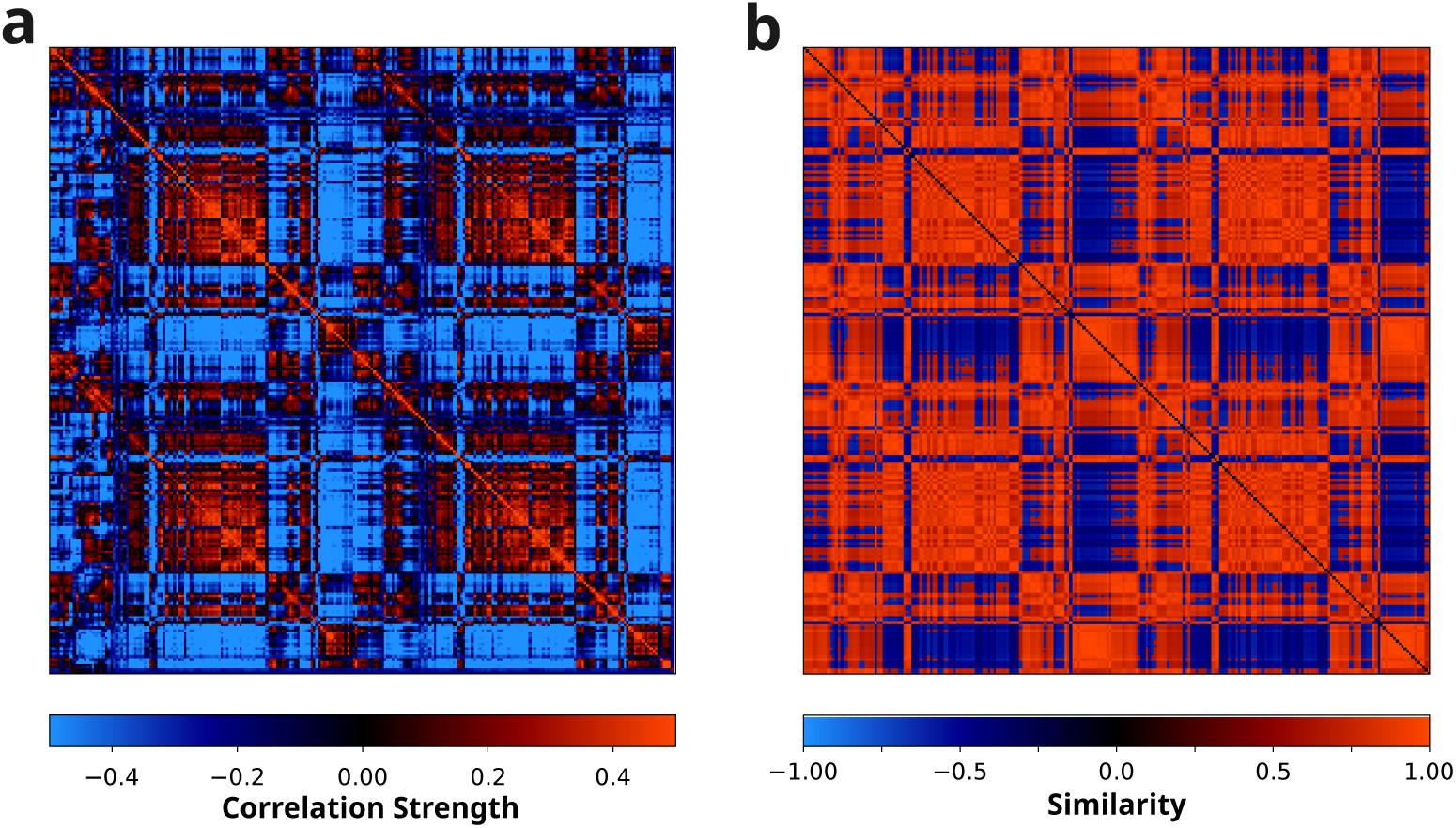
MeSiM Dimensionality Reduction. (a) Group-average MeSiM for the Geneva sample (*N* = 51). (b) Reconstructed MeSiM from PCA- and t-SNE–based dimensionality reduction of the group-average Geneva sample’s nodal connectivity profiles.

**Supplementary Fig. 3.**
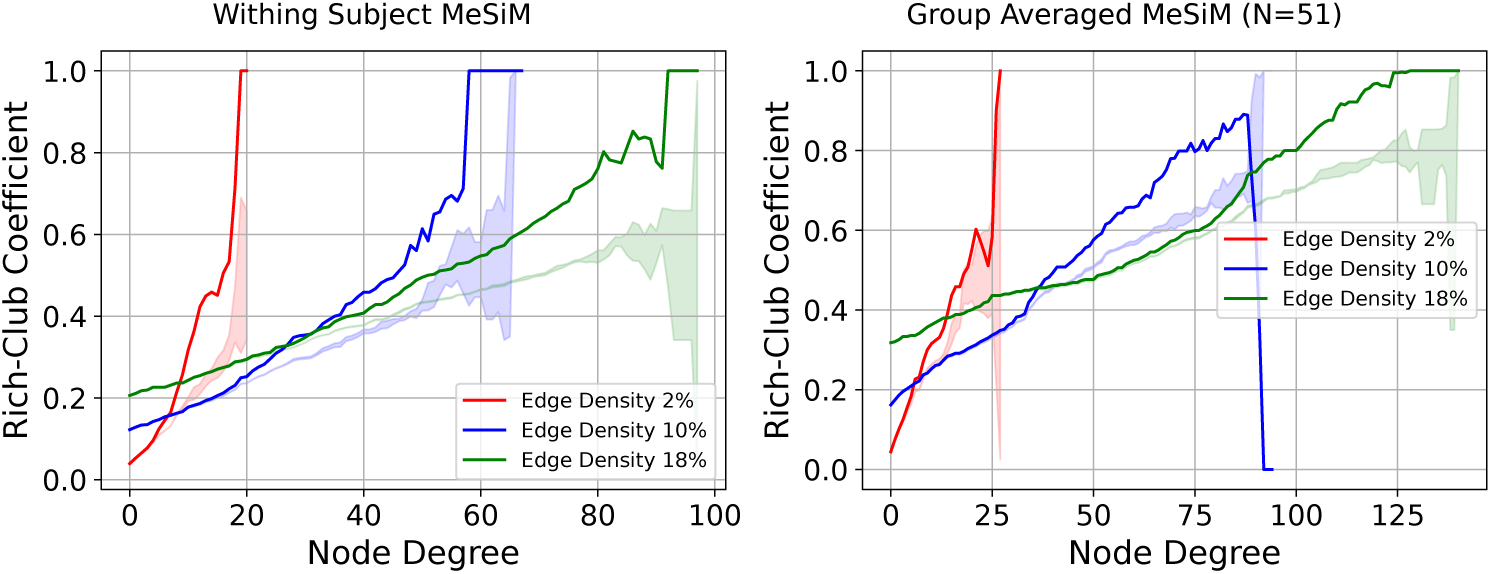
MeSiM Rich-Club Coefficient Analysis. Rich-club coefficients for MeSiM matrices compared to Erdős-Rényi random graphs. **a**, Rich-club coefficients for individual MeSiMs (*N* = 51) as a function of node degree at connection densities of 2%, 10%, and 18%. The shaded area comprises regions where the null hypothesis of absent rich club structure cannot be rejected (*α* = 0.05). **b**, Rich-club coefficients for the group-averaged MeSiM, with the same connection densities and random graph comparison.

**Supplementary Fig. 4.**
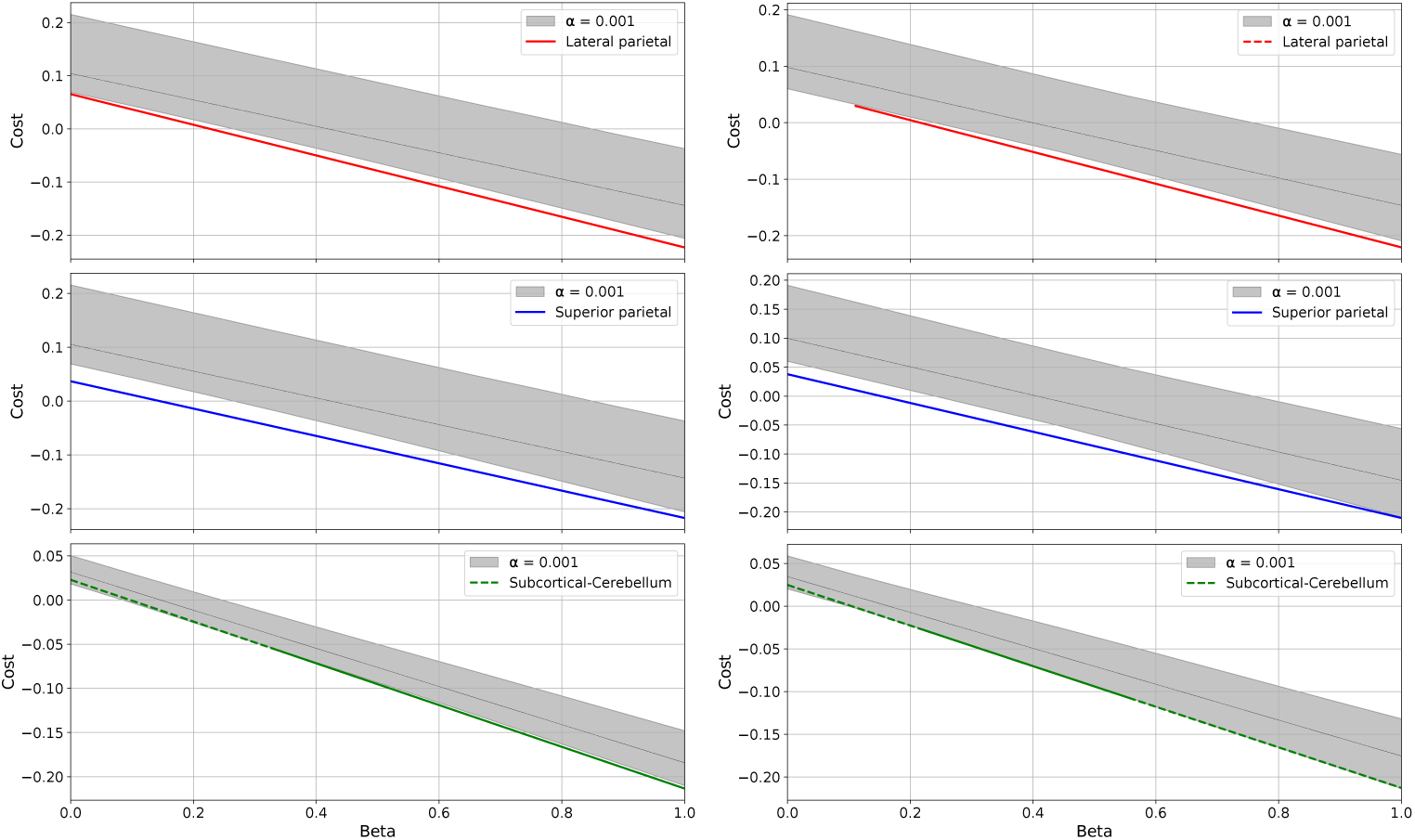
Metabolic Fibre Cost Function. Total cost functions are shown for both hemispheres (left side of the figure for the left hemisphere, right side for the right) across the three derived paths: the lateral parietal cortical path (top), the superior parietal cortical path (middle), and the subcortical-cerebellar path (bottom). Colored curves indicate the optimal solutions; shaded regions represent the other 99.999% of suboptimal solutions relative to the optimal one. The continuous curves mark the *β*-ranges over which the selected path’s cost function is statistically lower than that of the null distribution.

**Supplementary Fig. 5.**
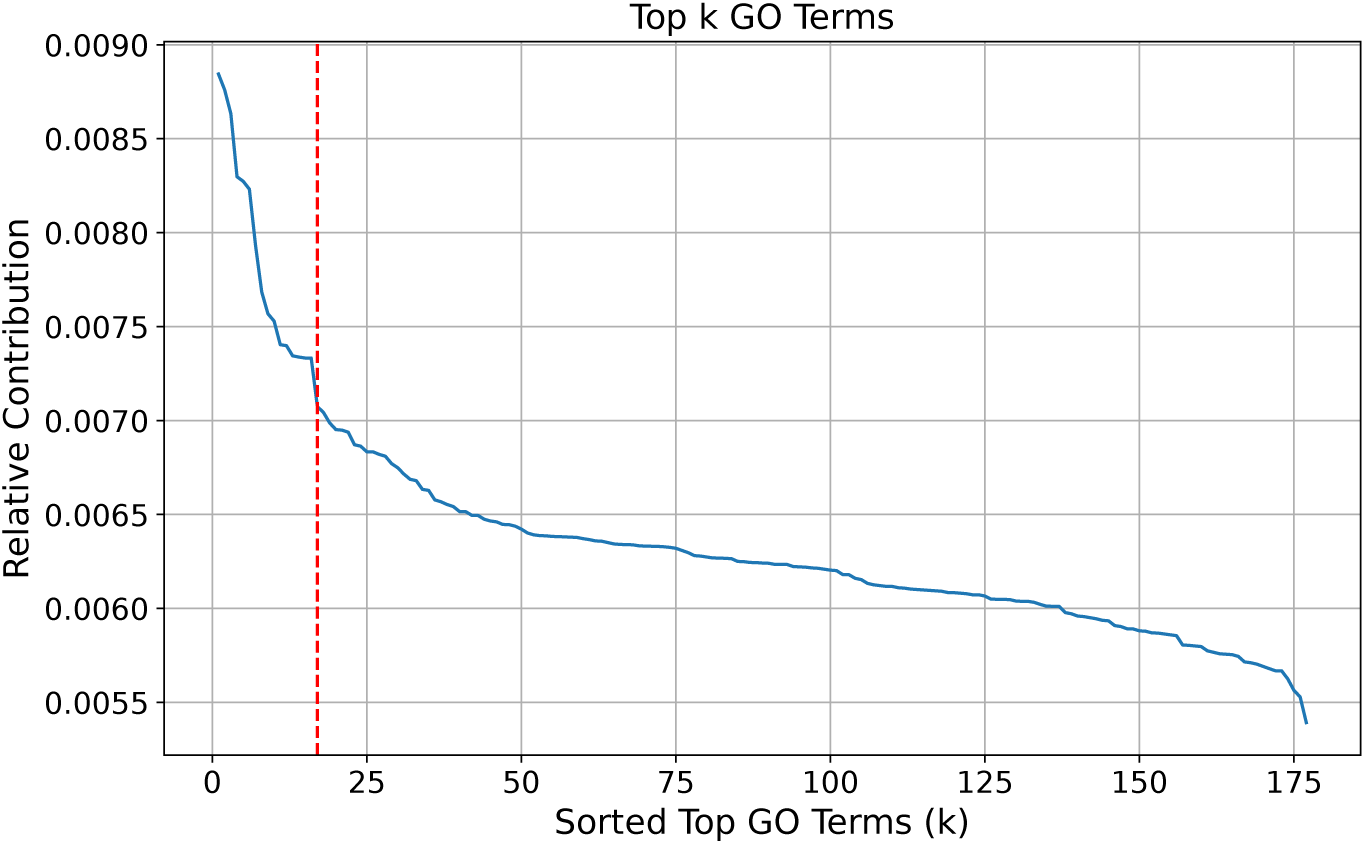
Leave-One-GO Analysis. Relative contribution was computed as the difference between the global edge-wise correlation of the group-average MeSiM (Geneva study) with the complete genetic co-expression matrix, and the correlation obtained when the genetic-coexpression matrix is imputed by the genetic expression profile corresponding to each left-out GO term. Contributions have ranked, and the elbow method identified 17 (vertical line) as the optimal number of GO terms contributing most to the overall correlation.

## References

1. Alemán-Gómez, Y. (2024). Chimera: An open source framework for combining multiple parcellations. https://github.com/connectomicslab/chimera. GitHub.

2. Alexander-Bloch, A.F., Shou, H., Liu, S., Satterthwaite, T.D., Glahn, D.C., Shinohara, R.T., … Raznahan, A. (2018). On testing for spatial correspondence between maps of human brain structure and function. Neuroimage, 178 , 540–551

3. Amunts, K., Lepage, C., Borgeat, L., Mohlberg, H., Dickscheid, T., Rousseau, M.-É., … others (2013). Bigbrain: an ultrahigh-resolution 3d human brain model. science, 340 (6139), 1472–1475

4. Barabási, A.-L. (2013). Network science. *Philosophical Transactions of the Royal Society A: Mathematical*, Physical and Engineering Sciences, 371 (1987), 20120375,

5. Baumann, P.S., Crespi, S., Marion-Veyron, R., Solida, A., Thonney, J., Favrod, J., … Conus, P. (2013). T reatment and e arly i ntervention in p sychosis p rogram (tipp-l ausanne): implementation of an early intervention programme for psychosis in s witzerland. Early intervention in psychiatry, 7 (3), 322–328,

6. Billot, B., Bocchetta, M., Todd, E., Dalca, A.V., Rohrer, J.D., Iglesias, J.E. (2020). Automated segmentation of the hypothalamus and associated subunits in brain mri. Neuroimage, 223 , 117287

7. Bullmore, E., & Sporns, O. (2009). Complex brain networks: graph theoretical analysis of structural and functional systems. Nature reviews neuroscience, 10 (3), 186–198

8. Bullmore, E., & Sporns, O. (2012). The economy of brain network organization. Nature reviews neuroscience, 13 (5), 336–349

9. Cammoun, L., Gigandet, X., Meskaldji, D., Thiran, J.P., Sporns, O., Do, K.Q., … Hagmann, P. (2012). Mapping the human connectome at multiple scales with diffusion spectrum mri. Journal of neuroscience methods, 203 (2), 386–397

10. Ceccarini, J., Liu, H., Van Laere, K., Morris, E.D., Sander, C.Y. (2020). Methods for quantifying neurotransmitter dynamics in the living brain with pet imaging. Frontiers in Physiology, 11, 792

11. Chen, Y., Wang, S., Hilgetag, C.C., Zhou, C. (2017). Features of spatial and functional segregation and integration of the primate connectome revealed by trade-off between wiring cost and efficiency. PLoS computational biology, 13 (9), e1005776

12. de Lange, S.C., Scholtens, L.H., Initiative, A.D.N., van den Berg, L.H., Boks, M.P., Bozzali, M., … others (2019). Shared vulnerability for connectome alterations across psychiatric and neurological brain disorders. Nature human behaviour , 3 (9), 988–998

13. de Reus, M.A., & van den Heuvel, M.P. (2013). Estimating false positives and negatives in brain networks. Neuroimage, 70 , 402–409

14. Estrada, E., & Hatano, N. (2008). Communicability in complex networks. *Physical Review E—Statistical*, Nonlinear, and Soft Matter Physics, 77 (3), 036111,

15. Finnema, S.J., Scheinin, M., Shahid, M., Lehto, J., Borroni, E., Bang-Andersen, B., … others (2015). Application of cross-species pet imaging to assess neurotransmitter release in brain. Psychopharmacology, 232, 4129–4157

16. Fischl, B., Salat, D.H., Busa, E., Albert, M., Dieterich, M., Haselgrove, C., … others (2002). Whole brain segmentation: automated labeling of neuroanatomical structures in the human brain. Neuron, 33 (3), 341–355

17. Friston, K. (2002). Beyond phrenology: what can neuroimaging tell us about distributed circuitry? Annual review of neuroscience, 25 (1), 221–250

18. Goulas, A., Betzel, R.F., Hilgetag, C.C. (2019). Spatiotemporal ontogeny of brain wiring. Science Advances, 5 (6), 10.1126/sciadv.aav9694

19. Guan, J., Yu, R., Wen, Y., Wu, H., Qin, H., Zhang, Q., Chen, W. (2017). Detection and application of neurochemical profile by multiple regional 1h-mrs in parkinson’s disease. Brain and Behavior , 7 (9), , 10.1002/brb3.792

20. Guo, J.U., Dengke, K., Mo, H., Ball, M.P., Jang, M.H., Bonaguidi, M.A., … Song, H. (2011). Neuronal activity modifies the dna methylation landscape in the adult brain. Nature Neuroscience, 14 (10), 1345–1351, 10.1038/nn.2900

21. Hagmann, P. (2005). From diffusion mri to brain connectomics (Unpublished doctoral dissertation). EPFL.

22. Hagmann, P., Cammoun, L., Gigandet, X., Meuli, R., Honey, C.J., Wedeen, V.J., Sporns, O. (2008). Mapping the structural core of human cerebral cortex. PLoS biology, 6 (7), e159,

23. Hansen, J.Y., Shafiei, G., Voigt, K., Liang, E.X., Cox, S.M., Leyton, M., … Misic, B. (2023). Integrating multimodal and multiscale connectivity blueprints of the human cerebral cortex in health and disease. PLoS biology, 21 (9), e3002314,

24. Hawrylycz, M.J., Lein, E.S., Guillozet-Bongaarts, A.L., Shen, E.H., Ng, L., Miller, J.A., … others (2012). An anatomically comprehensive atlas of the adult human brain transcriptome. Nature, 489 (7416), 391–399

25. Heinzle, J., & Stephan, K.E. (2018). Dynamic causal modeling and its application to psychiatric disorders. Computational psychiatry (pp. 117–144). Elsevier.

26. Horská, A., & Barker, P.B. (2010). Imaging of brain tumors: Mr spectroscopy and metabolic imaging. Neuroimaging Clinics, 20 (3), 293–310,

27. Iglesias, J.E., Augustinack, J.C., Nguyen, K., Player, C.M., Player, A., Wright, M., … others (2015). A computational atlas of the hippocampal formation using ex vivo, ultra-high resolution mri: Application to adaptive segmentation of in vivo mri. Neuroimage, 115 , 117–137

28. Klauser, A., Courvoisier, S., Kasten, J., Kocher, M., Guerquin-Kern, M., Van De Ville, D., Lazeyras, F. (2019). Fast high-resolution brain metabolite mapping on a clinical 3t mri by accelerated h-fid-mrsi and low-rank constrained reconstruction. Magnetic resonance in medicine, 81 (5), 2841–2857,

29. Klauser, A., Klauser, P., Grouiller, F., Courvoisier, S., Lazeyras, F. (2022). Wholebrain high-resolution metabolite mapping with 3d compressed-sensing sense lowrank 1h fid-mrsi. NMR in Biomedicine, 35 (1), e4615,

30. Klauser, A., Strasser, B., Thapa, B., Lazeyras, F., Andronesi, O. (2021). Achieving high-resolution 1h-mrsi of the human brain with compressed-sensing and lowrank reconstruction at 7 tesla. Journal of Magnetic Resonance, 331 , 107048,

31. Klopfenstein, D., Zhang, L., Pedersen, B., Ramírez, F., Warwick Vesztrocy, A., Naldi, A., et al. (n.d.). Goatools: a python library for gene ontology analyses. sci rep. 2018; 8: 10872.

32. Kobak, D., & Berens, P. (2019). The art of using t-sne for single-cell transcriptomics. Nature communications, 10 (1), 5416,

33. Kondor, R.I., & Lafferty, J. (2002). Diffusion kernels on graphs and other discrete structures. Proceedings of the 19th international conference on machine learning (Vol. 2002, pp. 315–322).

34. Lecocq, A., Le Fur, Y., Maudsley, A.A., Le Troter, A., Sheriff, S., Sabati, M., … others (2015). Whole-brain quantitative mapping of metabolites using short echo three-dimensional proton mrsi. Journal of Magnetic Resonance Imaging, 42 (2), 280–289

35. Linderman, G.C., & Steinerberger, S. (2019). Clustering with t-sne, provably. SIAM journal on mathematics of data science, 1 (2), 313–332

36. Ma, J., Chen, X., Gu, Y., Li, L., Cam-CAN, Lin, Y., Dai, Z. (2023). Trade-offs among cost, integration, and segregation in the human connectome. Network Neuroscience, 7 (2), 604–631

37. Martin, G. (2012). Network analysis and the connectopathies: current research and future approaches. Nonlinear dynamics, psychology, and life sciences, 16 (1), 79–90

38. Maudsley, A.A., Domenig, C., Govind, V., Darkazanli, A., Studholme, C., Arheart, K., Bloomer, C. (2009). Mapping of brain metabolite distributions by volumetric proton mr spectroscopic imaging (mrsi). Magnetic Resonance in Medicine: An Official Journal of the International Society for Magnetic Resonance in Medicine, 61 (3), 548–559

39. Moser, P., Eckstein, K., Hingerl, L., Weber, M., Motyka, S., Strasser, B., … Bogner, W. (2020). Intra-session and inter-subject variability of 3d-fid-mrsi using single-echo volumetric epi navigators at 3t. Magnetic resonance in medicine, 83 (6), 1920–1929,

40. Najdenovska, E., Alemán-Gómez, Y., Battistella, G., Descoteaux, M., Hagmann, P., Jacquemont, S., … Bach Cuadra, M. (2018). In-vivo probabilistic atlas of human thalamic nuclei based on diffusion-weighted magnetic resonance imaging. Scientific data, 5 (1), 1–11,

41. Newlin, N.R., Rheault, F., Schilling, K.G., Landman, B.A. (2023). Characterizing streamline count invariant graph measures of structural connectomes. Journal of Magnetic Resonance Imaging, 58 (4), 1211–1220,

42. Oldham, M.C., Horvath, S., Geschwind, D.H. (2006). Conservation and evolution of gene coexpression networks in human and chimpanzee brains. Proceedings of the National Academy of Sciences, 103 (47), 17973–17978,

43. Pichler, B.J., Wehrl, H.F., Kolb, A., Judenhofer, M.S. (2008). Positron emission tomography/magnetic resonance imaging: the next generation of multimodality imaging? Seminars in nuclear medicine (Vol. 38, pp. 199–208).

44. Piguet, C., Klauser, P., Celen, Z., James Murray, R., Magnus Smith, M., Merglen, A. (2022). Randomized controlled trial of a mindfulness-based intervention in adolescents from the general population: The mindfulteen neuroimaging study protocol. Early Intervention in Psychiatry, 16 (8), 891–901,

45. Pletikos, M., Sousa, A.M., Sedmak, G., Meyer, K.A., Zhu, Y., Cheng, F., … Šestan, N. (2014). Temporal specification and bilaterality of human neocortical topographic gene expression. Neuron, 81 (2), 321–332,

46. Preisig, M., Fenton, B.T., Matthey, M.-L., Berney, A., Ferrero, F. (1999). Diagnostic interview for genetic studies (digs): inter-rater and test-retest reliability of the french version. European archives of psychiatry and clinical neuroscience, 249, 174–179

47. Provencher, S.W. (2001). Automatic quantitation of localized in vivo 1h spectra with lcmodel. NMR in Biomedicine: An International Journal Devoted to the Development and Application of Magnetic Resonance In Vivo, 14 (4), 260–264

48. Ratai, E., Alshikho, M., Zürcher, N., Loggia, M., Cebulla, C., Cernasov, P., … Atassi, N. (2018). Integrated imaging of [11c]-pbr28 pet, mr diffusion and magnetic resonance spectroscopy 1h-mrs in amyotrophic lateral sclerosis. Neuroimage Clinical, 20 , 357–364, 10.1016/j.nicl.2018.08.007

49. Sander, C.Y., Hansen, H.D., Wey, H.-Y. (2020). Advances in simultaneous pet/mr for imaging neuroreceptor function. Journal of cerebral blood flow & metabolism, 40 (6), 1148–1166,

50. Saygin, Z.M., Kliemann, D., Iglesias, J.E., van der Kouwe, A.J., Boyd, E., Reuter, M., … others (2017). High-resolution magnetic resonance imaging reveals nuclei of the human amygdala: manual segmentation to automatic atlas. Neuroimage, 155, 370–382,

51. Schaefer, A., Kong, R., Gordon, E.M., Laumann, T.O., Zuo, X.-N., Holmes, A.J., … Yeo, B.T. (2018). Local-global parcellation of the human cerebral cortex from intrinsic functional connectivity mri. Cerebral cortex, 28 (9), 3095–3114

52. Scholtens, L.H., Schmidt, R., de Reus, M.A., van den Heuvel, M.P. (2014). Linking macroscale graph analytical organization to microscale neuroarchitectonics in the macaque connectome. Journal of Neuroscience, 34 (36), 12192–12205

53. Seghier, M.L., Zeidman, P., Neufeld, N.H., Leff, A.P., Price, C.J. (2010). Identifying abnormal connectivity in patients using dynamic causal modeling of fmri responses. Frontiers in systems neuroscience, 4 , 142

54. Seidlitz, J., Váša, F., Shinn, M., Romero-Garcia, R., Whitaker, K.J., Vértes, P.E., … others (2018). Morphometric similarity networks detect microscale cortical organization and predict inter-individual cognitive variation. Neuron, 97 (1), 231–247,

55. Shan, Y., Wang, Z., Song, S., Xue, Q., Ge, Q., Yang, H., … Lu, J. (2022). Integrated positron emission tomography/magnetic resonance imaging for resting-state functional and metabolic imaging in human brain: what is correlated and what is impacted. Frontiers in Neuroscience, 16 , 824152

56. Shin, J., Ming, G.L., Song, H. (2014). Dna modifications in the mammalian brain. Philosophical Transactions of the Royal Society B: Biological Sciences, 369 (1652), 20130512, 10.1098/rstb.2013.0512

57. Smith, R.E., Tournier, J.-D., Calamante, F., Connelly, A. (2013). Sift: Spherical-deconvolution informed filtering of tractograms. Neuroimage, 67 , 298–312

58. Solari, S.V.H., & Stoner, R.M. (2011). Cognitive consilience: primate non-primary neuroanatomical circuits underlying cognition. Frontiers in neuroanatomy, 5 , 13956

59. Sporns, O., Tononi, G., Kötter, R. (2005). The human connectome: a structural description of the human brain. PLoS computational biology, 1 (4), e42

60. Stam, C.J. (2014). Modern network science of neurological disorders. Nature Reviews Neuroscience, 15 (10), 683–695

61. Steel, A., Chiew, M., Jezzard, P., Voets, N.L., Plaha, P., Thomas, M.A., … Emir, U.E. (2018). Metabolite-cycled density-weighted concentric rings k-space trajectory (dw-crt) enables high-resolution 1 h magnetic resonance spectroscopic imaging at 3-tesla. Scientific reports, 8 (1), 7792

62. Stephan, K.E., Kasper, L., Harrison, L.M., Daunizeau, J., den Ouden, H.E., Breakspear, M., Friston, K.J. (2008). Nonlinear dynamic causal models for fmri. Neuroimage, 42 (2), 649–662

63. Steullet, P., Cabungcal, J., Monin, A., Dwir, D., O’donnell, P., Cuenod, M., Do, K. (2016). Redox dysregulation, neuroinflammation, and nmda receptor hypofunction: a “central hub” in schizophrenia pathophysiology? Schizophrenia research, 176 (1), 41–51

64. Stiernman, L.J., Grill, F., Hahn, A., Rischka, L., Lanzenberger, R., Panes Lundmark, V., … Rieckmann, A. (2021). Dissociations between glucose metabolism and blood oxygenation in the human default mode network revealed by simultaneous pet-fmri. Proceedings of the National Academy of Sciences, 118 (27), e2021913118

65. Tournier, J.-D., Calamante, F., Connelly, A. (2012). Mrtrix: diffusion tractography in crossing fiber regions. International journal of imaging systems and technology, 22 (1), 53–66

66. Tournier, J.-D., Smith, R., Raffelt, D., Tabbara, R., Dhollander, T., Pietsch, M., … Connelly, A. (2019). Mrtrix3: A fast, flexible and open software framework for medical image processing and visualisation. Neuroimage, 202 , 116137

67. Tustison, N.J., Cook, P.A., Holbrook, A.J., Johnson, H.J., Muschelli, J., Devenyi, G.A., … Avants, B.B. (2021, April). The ANTsX ecosystem for quantitative biological and medical imaging. Scientific Reports, 11 (1), 9068, 10.1038/s41598-021-87564-6 Retrieved from https://doi.org/10.1038/s41598-021-87564-6

68. Urchs, S., Armoza, J., Moreau, C., Benhajali, Y., St-Aubin, J., Orban, P., Bellec, P. (2019). Mist: A multi-resolution parcellation of functional brain networks. MNI Open Research, 1, 3

69. Vaishnavi, S.N., Vlassenko, A.G., Rundle, M.M., Snyder, A.Z., Mintun, M.A., Raichle, M.E. (2010). Regional aerobic glycolysis in the human brain. Proceedings of the National Academy of Sciences, 107 (41), 17757–17762

70. van den Heuvel, M.P., Scholtens, L.H., Barrett, L.F., Hilgetag, C.C., de Reus, M.A. (2015). Bridging cytoarchitectonics and connectomics in human cerebral cortex. Journal of Neuroscience, 35 (41), 13943–13948

71. van den Heuvel, M.P., & Sporns, O. (2019). A cross-disorder connectome landscape of brain dysconnectivity. Nature reviews neuroscience, 20 (7), 435–446,

72. von Békésy, G. (1970). Travelling waves as frequency analysers in the cochlea. Nature, 225 (5239), 1207–1209,

73. von Economo, C.F., & Koskinas, G.N. (1925). Die cytoarchitektonik der hirnrinde des erwachsenen menschen. J. Springer.

74. Wang, M., Yan, Z., Xiao, S.-y., Zuo, C., Jiang, J. (2020). A novel metabolic connectome method to predict progression to mild cognitive impairment. Behavioural Neurology, 2020 (1), 2825037

75. Wedeen, V.J., Hagmann, P., Tseng, W.-Y.I., Reese, T.G., Weisskoff, R.M. (2005). Mapping complex tissue architecture with diffusion spectrum magnetic resonance imaging. Magnetic resonance in medicine, 54 (6), 1377–1386,

76. Wei, Y., Scholtens, L., Turk, E., Van den Heuvel, M. (2019). Multiscale examination of cytoarchitectonic similarity and human brain connectivity. netw neurosci. 3: 124–137.

77. Wei, Y., Scholtens, L.H., Turk, E., Van Den Heuvel, M.P. (2018). Multiscale examination of cytoarchitectonic similarity and human brain connectivity. Network Neuroscience, 3 (1), 124–137,

78. Xia, M., Sun, X., Bu, X., Li, Q., He, Y. (2024). Big connectome imaging data in psychiatric disorders. Medicine Plus, 100038,

79. Yendiki, A., Koldewyn, K., Kakunoori, S., Kanwisher, N., Fischl, B. (2014). Spurious group differences due to head motion in a diffusion mri study. Neuroimage, 88, 79–90

80. Zalesky, A., Fornito, A., Cocchi, L., Gollo, L.L., van den Heuvel, M.P., Breakspear, M. (2016). Connectome sensitivity or specificity: which is more important? Neuroimage, 142, 407–420

81. Zimny, A., Szmyrka, M., Szewczyk, P., Bladowska, J., Pokryszko-Dragan, A., Gruszka, E., … Sąsiadek, M. (2013). In vivo evaluation of brain damage in the course of systemic lupus erythematosus using magnetic resonance spectroscopy, perfusion-weighted and diffusion-tensor imaging. Lupus, 23 (1), 10–19, 10.1177/0961203313511556

